# An attenuated vaccinia vaccine encoding the SARS-CoV-2 spike protein elicits broad and durable immune responses, and protects cynomolgus macaques and human ACE2 transgenic mice from SARS-CoV-2 and its variants

**DOI:** 10.1101/2022.06.12.495779

**Authors:** Hirohito Ishigaki, Fumihiko Yasui, Misako Nakayama, Akinori Endo, Naoki Yamamoto, Kenzaburo Yamaji, Cong Thanh Nguyen, Yoshinori Kitagawa, Takahiro Sanada, Tomoko Honda, Tsubasa Munakata, Masahiko Higa, Sakiko Toyama, Risa Kono, Asako Takagi, Yusuke Matsumoto, Kaori Hayashi, Masanori Shiohara, Koji Ishii, Yasushi Saeki, Yasushi Itoh, Michinori Kohara

**Author notes:** These authors contributed equally: Hirohito Ishigaki, Fumihiko Yasui, Misako Nakayama. Corresponding authors: Michinori Kohara,; and Yasushi Itoh.

## Abstract

As long as the coronavirus disease 2019 (COVID-19) pandemic continues, new variants of severe acute respiratory syndrome coronavirus 2 (SARS-CoV-2) with altered antigenicity will emerge. The development of vaccines that elicit robust, broad, and durable protection against SARS-CoV-2 variants is urgently needed. We have developed a vaccine (rDIs-S) consisting of the attenuated vaccinia virus DIs strain platform carrying the SARS-CoV-2 *S* gene. rDIs-S induced neutralizing antibody and T-lymphocyte responses in cynomolgus macaques and human angiotensin converting enzyme 2 (hACE2) transgenic mice, and showed broad protection against SARS-CoV-2 isolates ranging from the early-pandemic strain (WK-521) to the recent Omicron BA. 1 variant (TY38-839). Using a tandem mass tag (TMT) -based quantitative proteomic analysis of lung homogenates from hACE2 transgenic mice, we found that, among mice subjected to challenge infection with WK-521, vaccination with rDIs-S prevented protein expression related to the severe pathogenic effects of SARS-CoV-2 infection (tissue destruction, inflammation, coagulation, fibrosis, and angiogenesis) and restored protein expression related to immune responses (antigen presentation and cellular response to stress). Furthermore, long-term studies in mice showed that rDIs-S maintains S protein-specific antibody titers for at least 6 months after a 1^st^ vaccination. Thus, rDIs-S appears to provide broad and durable protective immunity against SARS-CoV-2, including current and possibly future variants.

## Introduction

Coronavirus disease 2019 (COVID-19), which is caused by infection with severe acute respiratory syndrome coronavirus 2 (SARS-CoV-2), has spread worldwide due to the lack of specific immunity against SARS-CoV-2 in most humans ^1,2^. Since the outbreak began in December 2019, SARS-CoV-2 infection has been associated with more than 510 million cases, resulting in more than 6.2 million deaths worldwide (https://covid19.who.int/). The acquisition of memory immune responses against SARS-CoV-2 is required for preventing COVID-19 and severe symptoms that require hospitalization. Vaccination is considered an essential means of obtaining such lymphocytic responses prior to infection.

SARS-CoV-2 is an enveloped single-stranded, positive-sense RNA virus. The spike (S) protein on the virion surface mediates SARS-CoV-2 entry into target cells through binding to the host cell receptor, angiotensin converting enzyme 2 (ACE2) ^3,4^. Consistent with this fact, the S protein is a major target of both neutralizing antibodies (nAbs) ^5,6^ and T cell responses to SARS-CoV-2-infected cells ^7,8^, indicating that the S protein is important as a vaccine component to elicit protective immunity against SARS-CoV-2.

The global effort to develop an effective vaccine enabled the distribution of the first COVID-19 vaccines within a year of the start of the pandemic and the initial identification of SARS-CoV-2. Subsequently, several COVID-19 vaccines have been approved for general or emergency use in multiple countries, including the United States, the United Kingdom, China, and Russia (https://extranet.who.int/pqweb/vaccines/covid-19-vaccines), and more than ten billion doses have been administered worldwide (https://coronavirus.jhu.edu/map.html). Currently, mRNA vaccines ^9,10^, adenovirus vector vaccines ^11–13^, and inactivated whole-virus vaccines ^14,15^ are in wide use, but the development of additional vaccines that are safe and effective is still of interest, especially given concerns about duration of protective efficacy, cross-reactivity against variants, vaccine cost, and the need for cold chains for distribution of the current vaccines.

Highly attenuated vaccinia viruses (VACVs) have gained attention as promising viral vectors owing to their safety and immunogenicity in humans, properties that have contributed to the eradication of smallpox ^16^. Among VACVs, the DIs strain was derived from the DIE strain of VACV through extensive serial passaging using one-day-old eggs ^17^. Notably, the DIs strain has a restricted host range because of a large-scale deletion (approximately 15.4 kb) representing 8% of the parental VACV genome; this deletion results in a loss of replication in most mammalian cells. The recombinant DIs strain also has been tested extensively as a platform for a candidate vaccine against severe acute respiratory syndrome (SARS), a previous coronavirus outbreak ^18,19^. Thus, the DIs strain is considered a promising viral vector for the development of novel vaccines. In the present study, we investigated the protective efficacy of rDIs-S, a recombinant DIs strain carrying the SARS-CoV-2 spike-encoding gene, against SARS-CoV-2 infection in both a nonhuman primate model and a human ACE2 (hACE2) transgenic mouse model.

## Results

### Immunization with rDIs-S induces both humoral and cellular immune responses, and protects hACE2 transgenic mice from lethal challenge with SARS-CoV-2 and its variants

rDIs-S was constructed by homologous gene recombination in primary chicken embryonic fibroblasts infected with DIs and transfected with the pSMART-DIs-L3 plasmid vector (Fig. 1A). This plasmid carries the full-length spike protein-encoding *S* gene of an early-pandemic SARS-CoV-2 strain (AI/I-004/2020 strain GISAID EPI_ISL_407084). Western blotting confirmed the expression of S protein in VeroE6/TMPRSS2 cells infected with rDIs-S (Fig. 1B). To determine the immunogenicity of rDIs-S, S protein-specific humoral and cellular immune responses were analyzed in C57BL/6 mice that had been immunized intradermally with either rDIs-S or DIs; immunization was performed two times with a 3-week interval between injections (Fig. 1C). In rDIs-S-inoculated mice, immunoglobulin (Ig) G specific for SARS-CoV-2 S protein and nAb were detected 1 week after the 1^st^ vaccination, and IgG and nAb levels increased after the 2^nd^ vaccination (Fig. 1D, E). In contrast, no S protein-specific antibodies were detected in the DIs-inoculated mice (control group). S protein-specific cellular immune responses were analyzed by *in vivo* cytotoxic T lymphocyte (CTL) assays. The *in vivo* number of target cells carrying SARS-CoV-2 S protein peptides was significantly reduced in the rDIs-S-inoculated mice compared to the DIs-inoculated mice (Fig. 1F).

**Fig. 1.**
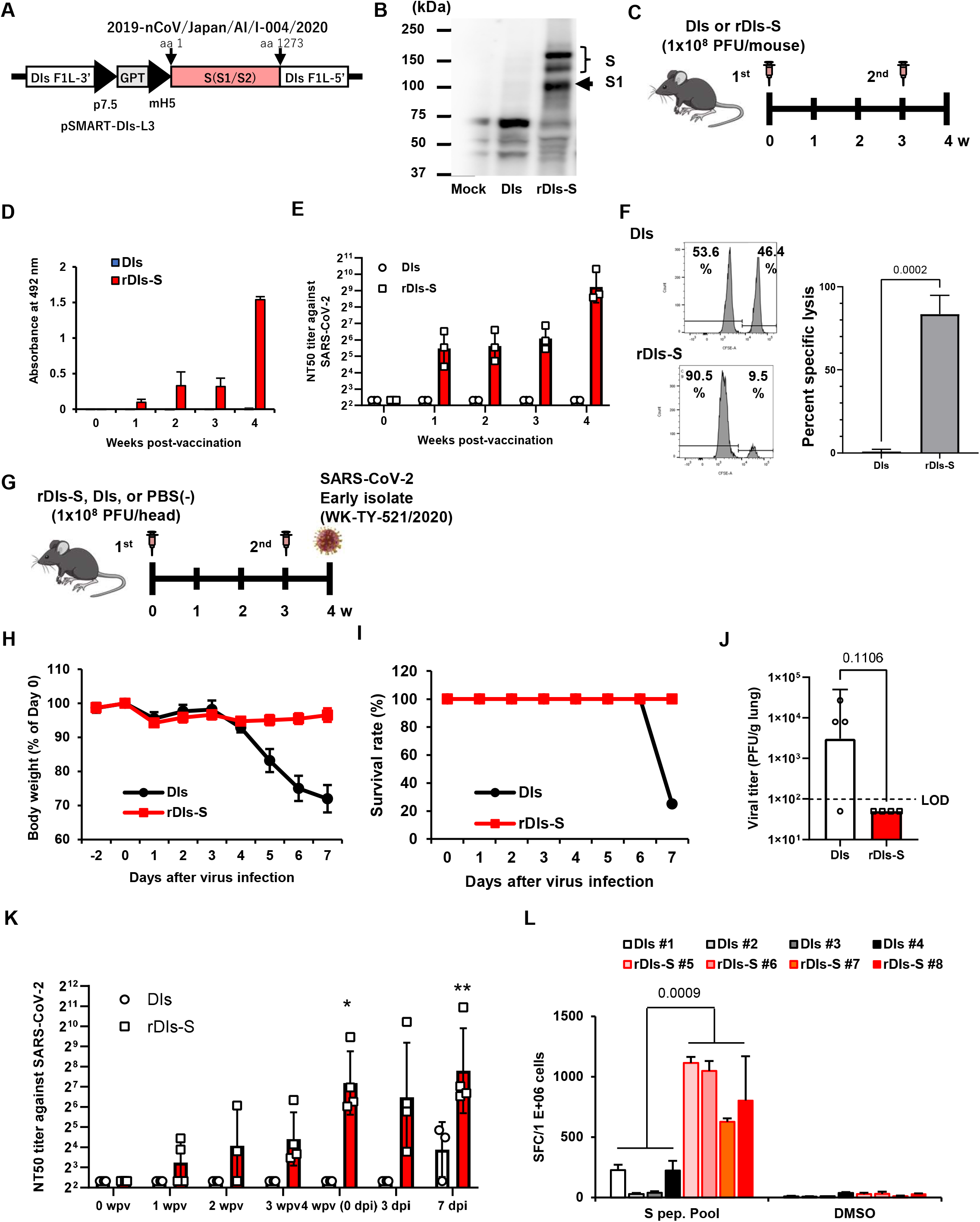
Immunization with rDIs-S induces both cellular and humoral immune responses. (A) Construction of the plasmid vector used for generating rDIs-S. (B) Expression of SARS-CoV-2 S protein as detected by western blot analysis. (C) Vaccination schedule in C57BL/6J mice. Nine to 10-week-old C57BL/6J mice were inoculated twice intradermally with 1 × 10^8^ PFU of rDIs-S or DIs at a 3-week interval. (D) Time course of the production of IgG specific for SARS-CoV-2 S protein as measured by ELISA (n = 3 per group). (E) Temporal changes in the neutralization titer against SARS-CoV-2 WK-521 strain (n = 3 per group). (F) An *in vivo* CTL assay specific for SARS-CoV-2 S protein peptides. The left panels are representative flow cytometry histograms. The right graph shows the mean ± SD of the specific killing of target cells (n = 3 per group). *p*-values were calculated using a two-tailed non-paired Student’s *t*-test. DIs: vaccinia virus DIs strain; rDIs-S: recombinant DIs carrying S gene of SARS-CoV-2; PFU: plaque forming unit; CTL: cytotoxic T lymphocyte; GPT: gene encoding xanthine-guanine phosphoribosyltransferase; p7.5: vaccinia virus early promoter; ELISA: enzyme-linked immunosorbent assay; hACE2: human angiotensin converting enzyme 2; SD: standard deviation. (G) Experimental schedule in hACE2 transgenic mice. Six to 10-week-old hACE2 transgenic mice were inoculated twice intradermally with 1 × 10^8^ PFU of rDIs-S or DIs at a 3-week interval, and then infected intratracheally with 20 TCID50 of SARS-CoV-2 (TY/WK-521/2020) 1 week after the 2^nd^ vaccination. (H-J) Protective effect of rDIs-S against challenge with SARS-CoV-2 WK-521 strain (n = 4 per group). (H) Temporal changes in the body weight of hACE2 transgenic mice with or without vaccination after infection with SARS-CoV-2 WK-521 strain. (I) Survival rate of hACE2 transgenic mice after SARS-CoV-2 infection. (J) Infectious viral titers in left lung homogenates were measured by a plaque assay. The dashed line indicates the limit of detection (LOD; 100 PFU/g lung). *p*-values were calculated using a two-tailed non-paired Student’s *t*-test. (K) Time-course of the neutralization titers against WK-521 before and after SARS-CoV-2 infection in hACE2 transgenic mice inoculated with either rDIs-S or DIs. *p*-values were calculated using a two-tailed Kruskal-Willis test, followed by a Dunn’s multiple comparison test (vs. 0 wpv). wpv: weeks post-vaccination; dpi: days post-infection; NT_50_: 50% neutralization titer; TCID_50_: 50% tissue culture infectious dose. (L) The number of IFN-γ-producing cells specific for SARS-CoV-2 S protein peptides examined by the ELISpot assay. *p*-values were calculated using a two-tailed non-paired Student’s *t*-test. IFN, interferon. SFC: spot forming cell.

Next, we examined the protective efficacy of rDIs-S against lethal challenge infection of hACE2 transgenic mice with an early-pandemic SARS-CoV-2 strain (Fig. 1G-J). The immunized hACE2 transgenic mice were infected intratracheally with SARS-CoV-2 (TY-WK-521/2020) 1 week after the 2^nd^ vaccination (Fig. 1G). All rDIs-S-inoculated mice survived the challenge with SARS-CoV-2 without any decrease in body weight, whereas DIs-inoculated mice succumbed to the SARS-CoV-2 infection, showing a drastic decrease in the body weight 4 days or more after the infection (Fig. 1H, I). When assessed at 7 days after infection, the titer of infectious SARS-CoV-2 titer in the lungs of rDIs-S-inoculated mice was below the detection limit; in contrast, the virus was detected in the lungs of 3 of 4 DIs-inoculated mice (Fig. 1J). In rDIs-S-inoculated hACE2 transgenic mice, nAb was detected 1 week after the 1^st^ vaccination, and nAb level increased after the 2^nd^ vaccination (Fig. 1K). The number of T cells specifically producing interferon (IFN) -γ was elevated significantly in the rDIs-S-inoculated group compared to the DIs-inoculated group (Fig. 1L).

As a next step, the cross-protective efficacy of rDIs-S against variants, including Beta, Delta, and Omicron BA.1, were investigated. The rDIs-S-inoculated hACE2 transgenic mice were infected intranasally with the Beta or Delta variant 1 week after the 2^nd^ vaccination (Fig. 2A). All rDIs-S-inoculated hACE2 mice, but not control (phosphate-buffered saline (PBS) -immunized) mice, survived the lethal challenge with the Beta variant of SARS-CoV-2 (TY8-612 strain) without any decrease in body weight (Fig. 2B, C). At 7 days after infection, the infectious SARS-CoV-2 titer in the lungs of rDIs-S-inoculated mice was below the detection limit in 3 of 4 mice, whereas the virus was detected in the lungs of all unvaccinated mice. Vaccination with rDIs-S also protected mice from lethal challenge with a Delta variant (TY11-927; Fig. 2E, F).

**Fig. 2.**
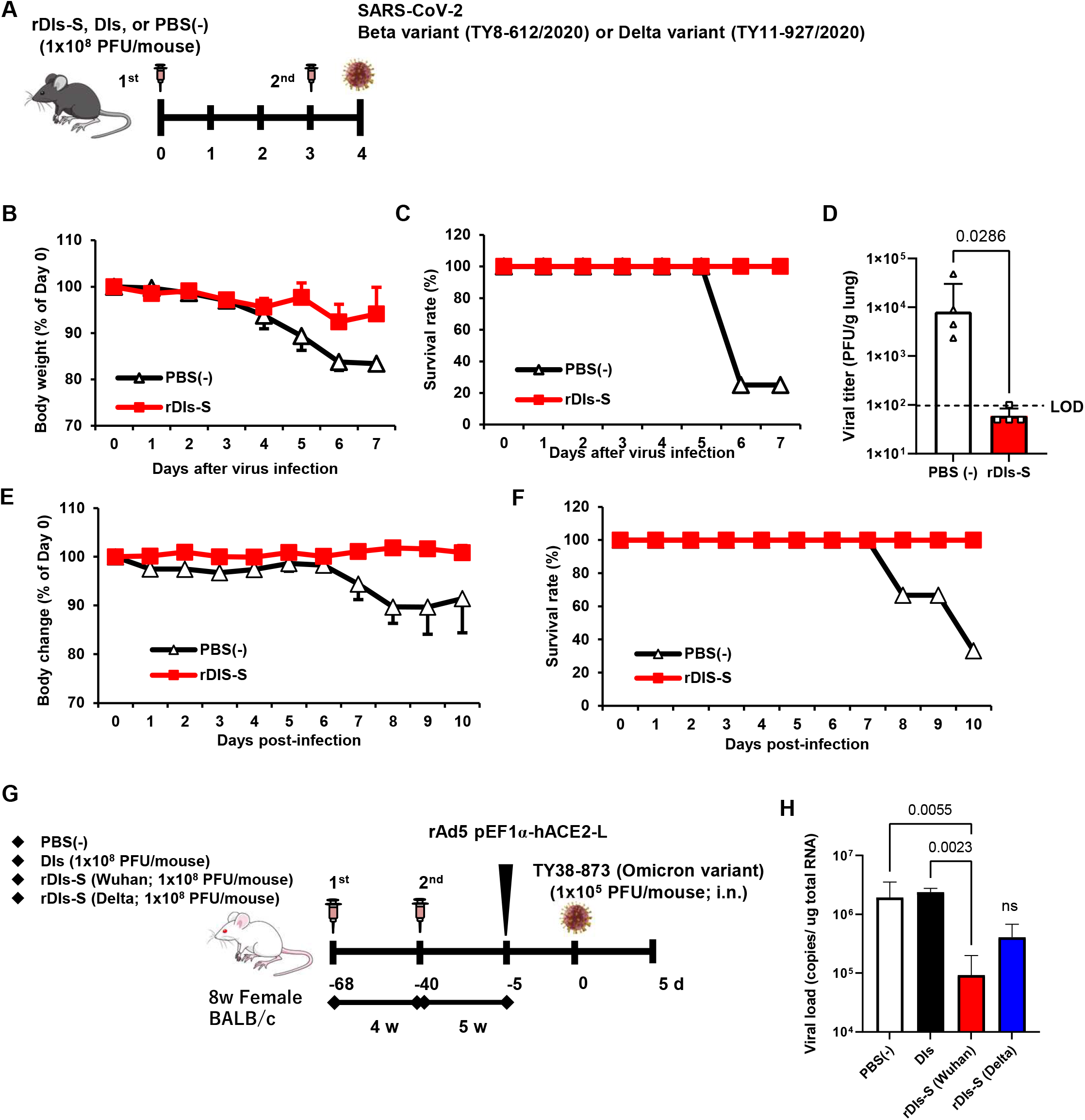
Immunization with rDIs-S protects mice from lethal challenge with SARS-CoV-2. (A) Experimental schedule in human angiotensin converting enzyme 2 (hACE2) transgenic mice. hACE2 transgenic mice were inoculated twice intradermally with 1 × 10^8^ PFU of rDIs-S or PBS(-) at a 3-week interval, and then infected intratracheally with 100 PFU of a Beta variant (TY8-612/2020) or 50 PFU of a Delta variant (TY11-927) SARS-CoV-2 strain 1 week after the 2^nd^ vaccination. (B-D) Protective effect of rDIs-S against challenge with Beta variant of SARS-CoV-2 (TY8-612 strain) (n = 4 per group). (B) Temporal changes in the body weight of hACE2 transgenic mice with or without vaccination after infection with Beta variant. (C) Survival rate of hACE2 transgenic mice after Beta variant infection. (D) Infectious viral titers in left lung homogenates were measured by a plaque assay. The dashed line indicates the limit of detection (LOD; 100 PFU/g lung). *p*-values were calculated using a two-tailed non-paired Student’s *t*-test. (E and F) Protective effect of rDIs-S against challenge with a Delta variant of SARS-CoV-2 (TY11-927 strain) (n = 3-4 per group). (E) Temporal changes in the body weight of hACE2 transgenic mice after infection with the TY11-927 strain. (F) Survival rates after infection with the TY11-927 strain. (G) Experimental schedule in BALB/c mice. BALB/c mice were inoculated twice epidermally with 1 × 10^8^ PFU of rDIs-S (derived from an early-pandemic strain of SARS-CoV-2 or a Delta variant), DIs, or PBS(-) at a 4-week interval. Five weeks after the 2^nd^ vaccination, the mice were intranasally inoculated with 5 × 10^7^ FFU of rAd5-pEF1a-hACE2 and challenged with 1 × 10^5^ PFU of an Omicron variant of SARS-CoV-2 five days later. (H) Viral RNA in left lung homogenates was quantified by qRT-PCR. *p*-values were calculated using a two-tailed Kruskal-Willis test, followed by a Dunn’s multiple comparison test. ns: not significant; FFU: focus forming unit; PFU: plaque forming unit; PBS: phosphate buffered saline.

As we reported elsewhere ^20^, we recently generated an adenoviral vector expressing the hACE2-encoding gene under control of the *EF1α* promoter with a leftward orientation (rAd5 pEF1α-hACE2-L); this novel transgene vector confers SARS-CoV-2 susceptibility in wild-type mice. Using this model, we investigated the ability of rDIs-S carrying the *S* gene from an early-pandemic strain of SARS-CoV-2 or from a Delta variant, to provide protection against the Omicron BA. 1 variant (TY38-839; Fig. 2G). BALB/c mice were inoculated twice (at a 4-week interval) by epicutaneous immunization (skin scarification) with rDIs-S encoding spike protein from either an early-pandemic strain of SARS-CoV-2 or a Delta variant. Five weeks after the 2^nd^ vaccination, the mice were inoculated intranasally with 5 × 10^7^ focus-forming units (FFU) of rAd5-pEF1a-hACE2 and challenged (5 days later) with the TY38-873 strain of SARS-CoV-2. Importantly, immunization with rDIs-S, which encodes spike derived from an early-pandemic strain of SARS-CoV-2, prevented the propagation of the virus, with an efficacy greater than that of rDIs-S encoding spike derived from a Delta variant (Fig. 2H). Taken together, these results indicated that rDIs-S efficiently protects mice from challenge not only with “classic” SARS-CoV-2 but also with viral variants. This protection is mediated by the induction of both humoral and cellular immune responses that are specific for the S protein of SARS-CoV-2.

### Protection from SARS-CoV-2 pneumonia in a nonhuman primate model vaccinated with rDIs-S

hACE2 transgenic mice infected with SARS-CoV-2 show much more severe acute morbidity than do human patients. Therefore, we next evaluated the efficacy of rDIs-S in a nonhuman primate model ^21^. Cynomolgus macaques were immunized intracutaneously with rDIs-S or DIs, administered twice with a 3-week interval between injections. To evaluate the protective efficacy of rDIs-S, the immunized macaques were infected with SARS-CoV-2 TY/WK-521/2020 via the conjunctiva, nasal cavity, oral cavity, and trachea 1 week after the 2^nd^ vaccination. Of the four macaques immunized with DIs, two and three macaques displayed infectious virus in nasal swab samples and lung tissue samples (respectively) at Day 7 after SARS-CoV-2 inoculation; in contrast, no infectious virus was detected in macaques immunized with rDIs-S, either in nasal swab samples at Day 3 or in lung tissue samples at Day 7 (Fig. 3A, B, Supplemental Tables 1, 2). Viral RNA was detected in the nasal swab samples, oral swab samples, and lung tissues of all four of the DIs-immunized macaques at 7 days after infection with SARS-CoV-2 virus (Fig. 3C, Supplemental Fig. 1). In contrast, in macaques immunized with rDIs-S, the levels of SARS-CoV-2 viral RNA in the trachea, bronchus, and a part of the lung tissues were below the detection limit. Thus, vaccination with rDIs-S prevented the propagation of SARS-CoV-2 in cynomolgus macaques.

**Fig. 3.**
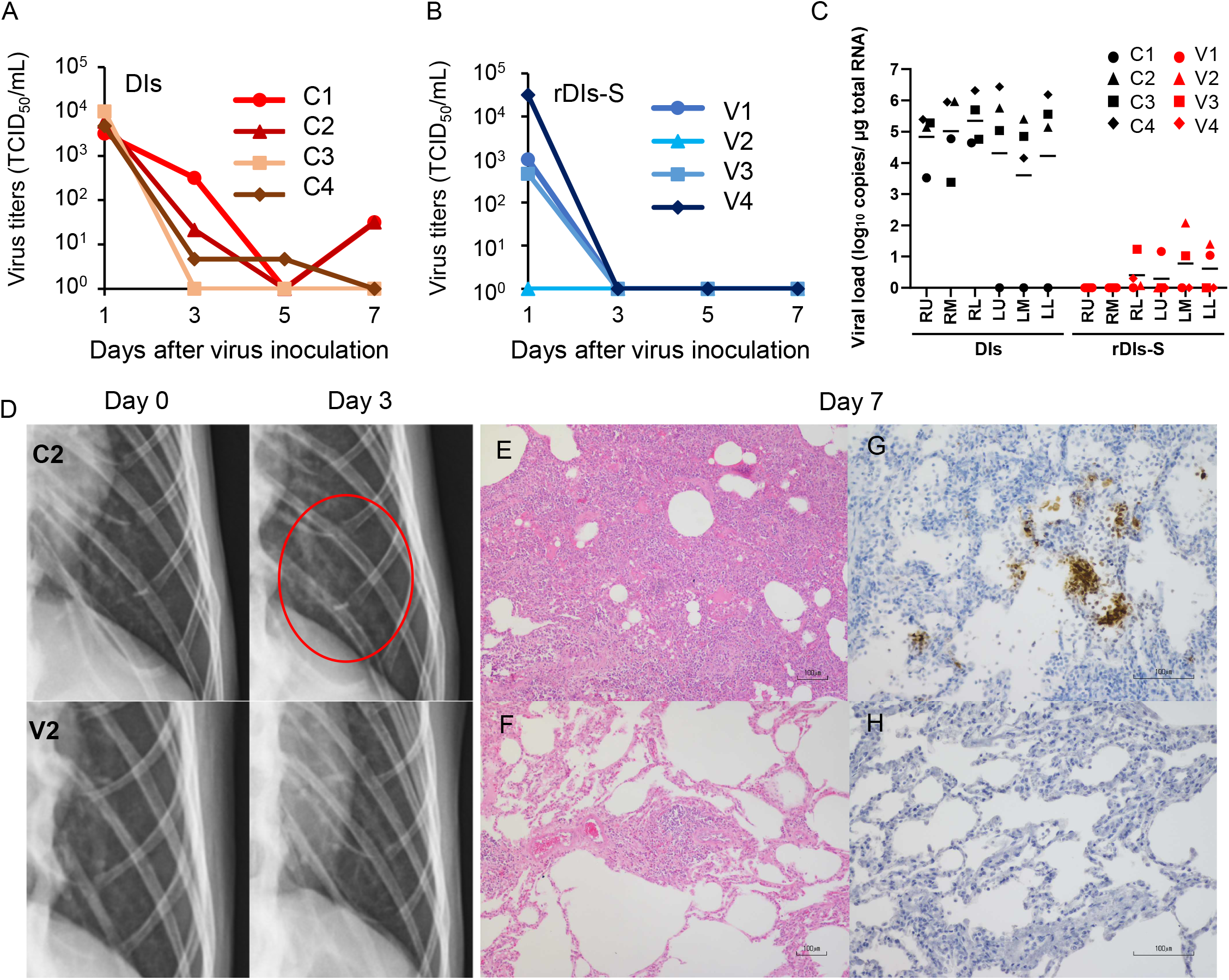
Protection from SARS-CoV-2 pneumonia in macaques vaccinated with rDIs-S. Cynomolgus macaques were immunized intradermally with DIs (C1 – C4) or rDIs-S (V1 – V4). One week after the 2^nd^ vaccination, SARS-CoV-2 WK-521 was inoculated into the conjunctiva, nostril, oral cavity, and trachea of each macaque on Day 0. (A, B) Nasal swab samples were collected on the indicated days after virus inoculation. Viral titers below the limit of detection (LOD; 0.67 log_10_TCID_50_/mL) are shown as 10^0^. (C) Lung tissues were collected 7 days after virus inoculation. Viral RNA was quantified by qRT-PCR. RU: right upper lobe; RM: right middle lobe; RL: right lower lobe; LU: left upper lobe; LM: left middle lobe; LL: left lower lobe. (D) X-ray radiography was performed on Day 0 before infection and on Day 3 after infection. Representative photos are shown. The red circle indicates the ground glass appearance. (E, F) Hematoxylin- and eosin-stained sections of lung. (G, H) Immunohistochemical staining of lung sections for SARS-CoV-2 N antigen (brown staining). Nuclei were stained with hematoxylin. Bars indicate 100 μm in E-H. (E, G) C2: a macaque immunized with DIs. (F, H) V2: a macaque immunized with rDIs-S. Bars indicate 100 μm.

We also examined the vaccinated macaques for clinical signs of disease after infection. All four of the DIs-immunized macaques showed increases in body temperature after infection with SARS-CoV-2 (Supplemental Fig. 2). Among these four macaques, three showed a body temperature higher than 39 °C during the daytime. In contrast, of four rDIs-S-immunized macaques (V1-V4), three did not show an increase in body temperature during the daytime, and only one (V4) showed a body temperature increase for the first 3 days after infection. Among the infected macaques, the clinical scores (which were determined based on body temperature, appetite, posture, and behavior (Supplemental Table 3)) were lower in the animals immunized with rDIs-S than in those immunized with DIs (Supplemental Fig. 3A-D). Thus, vaccination with rDIs-S attenuated the clinical signs of disease after SARS-CoV-2 infection in cynomolgus macaques.

Next, the effects of vaccination with rDIs-S on viral pneumonia were examined by X-ray radiography and histological examination of post-mortem samples. On chest X-ray radiography, all four macaques immunized with DIs showed a ground glass appearance in areas of the lungs by Day 3 after infection with SARS-CoV-2 (Fig. 3D). However, no apparent radiographic changes were detected in the lungs of macaques immunized with rDIs-S and subsequently infected with SARS-CoV-2. Macroscopic observations at necropsy revealed dark red lesions on the surfaces of the lungs in the macaques immunized with DIs and subsequently infected (Supplemental Fig. 4A), whereas a very mild reddish change was seen in the lung of only one of the four macaques (V4) vaccinated with rDIs-S and subsequently infected (Supplemental Fig. 4B). In macaques immunized with DIs, thickened alveolar walls, exudates, and hyaline membrane formation were observed in the lung tissues 7 days after infection with SARS-CoV-2; these changes were attenuated in the macaques immunized with rDIs-S and subsequently infected, as confirmed by the histological scoring (Fig. 3E, F, Supplemental Fig. 4C). At necropsy of the infected animals, the relative (body weight-normalized) lung weight of the macaques immunized with rDIs-S was significantly lower than that of the macaques immunized with DIs (Supplemental Fig. 4E), consistent with histological observations indicating pneumonia (Supplemental Fig. 4D). Additionally, in the infected animals, SARS-CoV-2 nucleoprotein (N protein) -positive cells formed clusters that were distributed sparsely in the lungs of the macaques immunized with DIs, whereas no N protein-positive cells were detected in the lung tissues of the macaques immunized with rDIs-S (Fig. 3G, H). Thus, vaccination with rDIs-S prevented viral pneumonia in cynomolgus macaques.

### Immune responses in a nonhuman primate model following vaccination with rDIs-S

We examined the acquired immune responses responsible for the protection in macaques immunized with rDIs-S. Among macaques immunized with rDIs-S, IgG antibodies specific for the SARS-CoV-2 S protein, including those with specificity for the receptor-binding domain (RBD) and domains 1 and 2 of the S protein (S1 and S2), were detected in the plasma 10 days after the 1^st^ vaccination (Fig. 4A, C), and the levels of those antibodies increased after the 2^nd^ vaccination. SARS-CoV-2-specific nAbs against the challenge strain WK-521 (Clade S) and variant strains SUMS2 (Clade GR, Pango lineage B1.1), QHN001 (Clade GRY, Pango lineage B.1.1.7), TY7-501 (Clade GR/501Y.V3, Pango lineage P. 1), and TY8-612 (Clade GH/501Y.v2, Pango lineage B. 1.351) (Supplemental Table 4) were detected in the plasma of the macaques immunized with rDIs-S, and the neutralization titers increased after challenge infection with WK-521, indicating the activation of memory responses after infection (Fig. 4B, D). On the other hand, no nAb specific for SARS-CoV-2 was detected in the plasma of the DIs-immunized macaques at Day 7 after challenge infection. T-cell responses specific for SARS-CoV-2 S protein peptides were detected 7 days after the 2^nd^ vaccination with rDIs-S (Fig. 4E, F). The ratio of IFN-γ- and interleukin (IL) −2-producing cells increased after the 2^nd^ rDIs-S vaccination and challenge infection. Thus, humoral and cellular immunity specific for SARS-CoV-2 was induced effectively in macaques immunized with rDIs-S.

**Fig. 4.**
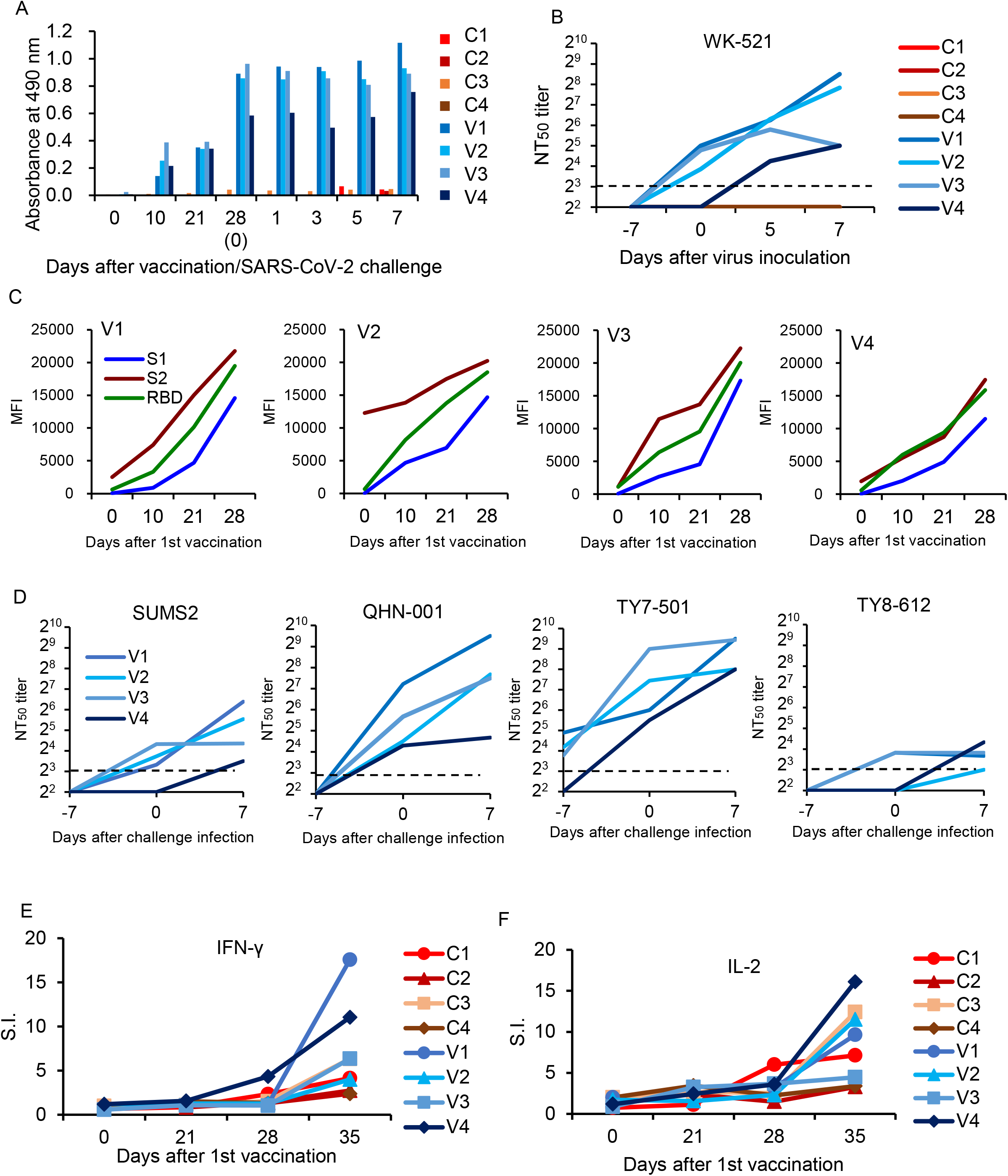
Immune responses in macaques vaccinated with rDIs-S. The macaques were immunized intradermally with DIs (C1 – C4) or rDIs-S (V1 – V4). One week after the 2^nd^ vaccination, SARS-CoV-2 WK-521 was inoculated on Day 0. (A) The levels of IgG specific for SARS-CoV-2 S protein in the plasma of macaques immunized with DIs (C1 – C4) and macaques immunized with rDIs-S (V1 – V4) were analyzed using ELISA. Plasma was collected after the 1^st^ vaccination (Days 0 to 28) and after challenge infection (Days 1 to 7). (B) 50% neutralization titers (NT50) of plasma against WK-521. Plasma was collected from macaques on the indicated days after the 1^st^ vaccination. Day -7: the day of the 2^nd^ vaccination and 7 days before challenge infection. Day 0: the day of challenge infection. Day-0 samples were collected before infection. Days 5 and 7: 5 and 7 days (respectively) after challenge infection. (C) Plasma IgG specific for SARS-CoV-2 S1, S2, and receptor-binding domain (RBD). MFI: mean fluorescence intensity. (D) Neutralizing antibody against variants of SARS-CoV-2. NT_50_ of plasma against variant strains with amino acid changes in the S protein were measured. (E and F) The numbers of IFN-γ-producing cells (E) and IL-2-producing cells (F) specific for SARS-CoV-2 S protein peptides were examined by the ELISpot assay. Stimulation indices (S.I.) were calculated as follows: S.I. = number of spots in the culture of cells with peptides / number of spots in the culture of cells without peptides.

### Prevention of inflammatory responses in macaques and hACE2 transgenic mice vaccinated with rDIs-S

The levels of systemic and local inflammation after SARS-CoV-2 infection and the effects of vaccination on cytokine responses were examined in macaques and hACE2 transgenic mice. The levels of the inflammatory cytokine IL-6 and the chemokine monocyte chemoattractant protein-1 (MCP-1) in the plasma of the macaques inoculated with DIs were increased on Day 1 after challenge infection, whereas no such increase was seen in the plasma of the macaques vaccinated with rDIs-S and subjected to challenge infection (Supplemental Fig. 5). The levels of IL-15 and granulocyte colony-stimulating factor (G-CSF) in the plasma of the macaques immunized with DIs showed a similar increase on Day 1 after challenge infection, and the apparent slight elevation persisted on Days 3 and 5. On the other hand, no significant increase in IL-15 or G-CSF levels was detected following challenge infection in the rDIs-S-vaccinated macaques. Thus, vaccination with rDIs-S prevented inflammatory responses in cynomolgus macaques.

Changes in protein expression levels in the lungs of hACE2 transgenic mice 7 days after virus infection were analyzed comprehensively by mass spectrometry (MS) -based quantitative proteomics using a tandem mass tag (TMT) reagent (Supplemental Fig. 6A). In the lungs of DIs-immunized hACE2 transgenic mice infected with WK-521, a total of 177 proteins showed significantly increased expression (mean fold-change ≥2.0, *p*-value <0.05), and 251 proteins showed significantly decreased expression (mean fold-change ≤0.5, *p*-value <0.05) compared to lung tissue from uninfected mice (Fig. 5A, top). On the other hand, the expression levels of 278 and 32 proteins were increased and decreased, respectively, in the lungs of infected mice that had been vaccinated with rDIs-S, compared to the uninfected mice (Fig. 5A, middle). The expression levels of 36 and 82 proteins were increased and decreased, respectively, in the lungs of infected mice that had been vaccinated with rDIs-S, compared to the infected mice that had been immunized with DIs (Fig. 5A, bottom). The proteins with increased and decreased levels in the DIs-immunized mice following infection were submitted for gene ontology (GO) term enrichment analyses using Metascape for terms related to biological processes (BP) ^22^. The proteins with increased levels in the infected mice that had been immunized with DIs were significantly enriched in terms related to phagocytosis, blood coagulation, and inflammatory response (Fig. 5B, upper), consistent with results obtained for COVID-19 patients. On the other hand, the proteins with decreased levels in the infected mice that had been immunized with DIs were significantly enriched in terms related to cytoplasmic translation, ribosome biogenesis, and negative regulation of chromatin silencing (Fig. 5B, lower), indicating that the *de novo* synthesis of proteins was significantly suppressed by SARS-CoV-2 infection. Of the 177 proteins with increased levels in DIs-immunized mice following infection, 57 showed decreases of greater than two-fold in rDIs-S-vaccinated mice following infection (Fig. 5A, bottom and Fig. 5C). In comparison, of the 251 proteins that were depleted in the DIs-immunized mice following infection, 28 showed increases of greater than 2-fold in the rDIs-S-vaccinated mice following infection (Fig. 5A, bottom and Fig. 5C). GO enrichment terms related to BP were analyzed for the 57 and 28 proteins that showed divergent changes in expression between the infected DIs-immunized mice and infected rDIs-S-vaccinated mice (Fig. 5D). All of the proteins listed under the top-13 GO enrichment terms of the upregulated proteins are shown in Supplemental Fig. 6B. Furthermore, overlap analysis of the proteins listed under the top-13 GO enrichment terms showed that these proteins include multiple GO enrichment terms (Supplemental Fig. 6C). The expression levels of proteins associated with fibrinolysis (coagulation), inflammatory proteins, and collagen catabolism (tissue destruction) were significantly lower in the rDIs-S-vaccinated mice than in the DIs-immunized mice, as were the expression levels of proteins related to leukocyte migration involved in inflammatory response, peptidase activity, defense responses to fungus, apoptotic signaling, responses to metal ions, smallmolecule biosynthesis processes, angiogenesis, responses to peptides, α-amino acid metabolic processes, and aminoglycan metabolic processes (Fig. 5E, Supplemental Fig. 6D). On the other hand, the expression levels of proteins involved in antigen presentation, negative regulation of cytokine production, chemotaxis, osteoblast differentiation, and cellular response to stress were decreased in the DIs-immunized mice compared to the rDIs-S-vaccinated mice and uninfected mice (Fig. 5F, Supplemental Fig. 7A to C). Although the expression levels of proteins involved in phagocytosis, oxidative stress, and protein transport were increased in the infected animals (whether DIs-immunized and rDIs-S-vaccinated; Supplemental Fig. 8), the expression levels tended to be lower in the rDIs-S-vaccinated mice than in the DIs-immunized mice. The expression levels of proteins involved in tissue repair processes, such as gene expression, cell-matrix adhesion, organelle organization, and cell part morphogenesis, were decreased even in the rDIs-S-vaccinated mice (Supplemental Fig. 9), but the number of proteins in this category was only 14. The magnitude of the decrease in the levels of these proteins was smaller in rDIs-S-vaccinated mice than in DIs-immunized mice. Taken together, these results indicated that, among mice subjected to challenge infection with WK-521, vaccination with rDIs-S prevents gene expression indicative of tissue destruction and of lung inflammation, and restores gene expression indicative of immune responses and tissue repair processes, changes that are otherwise observed in DIs-immunized mice upon infection.

**Fig. 5.**
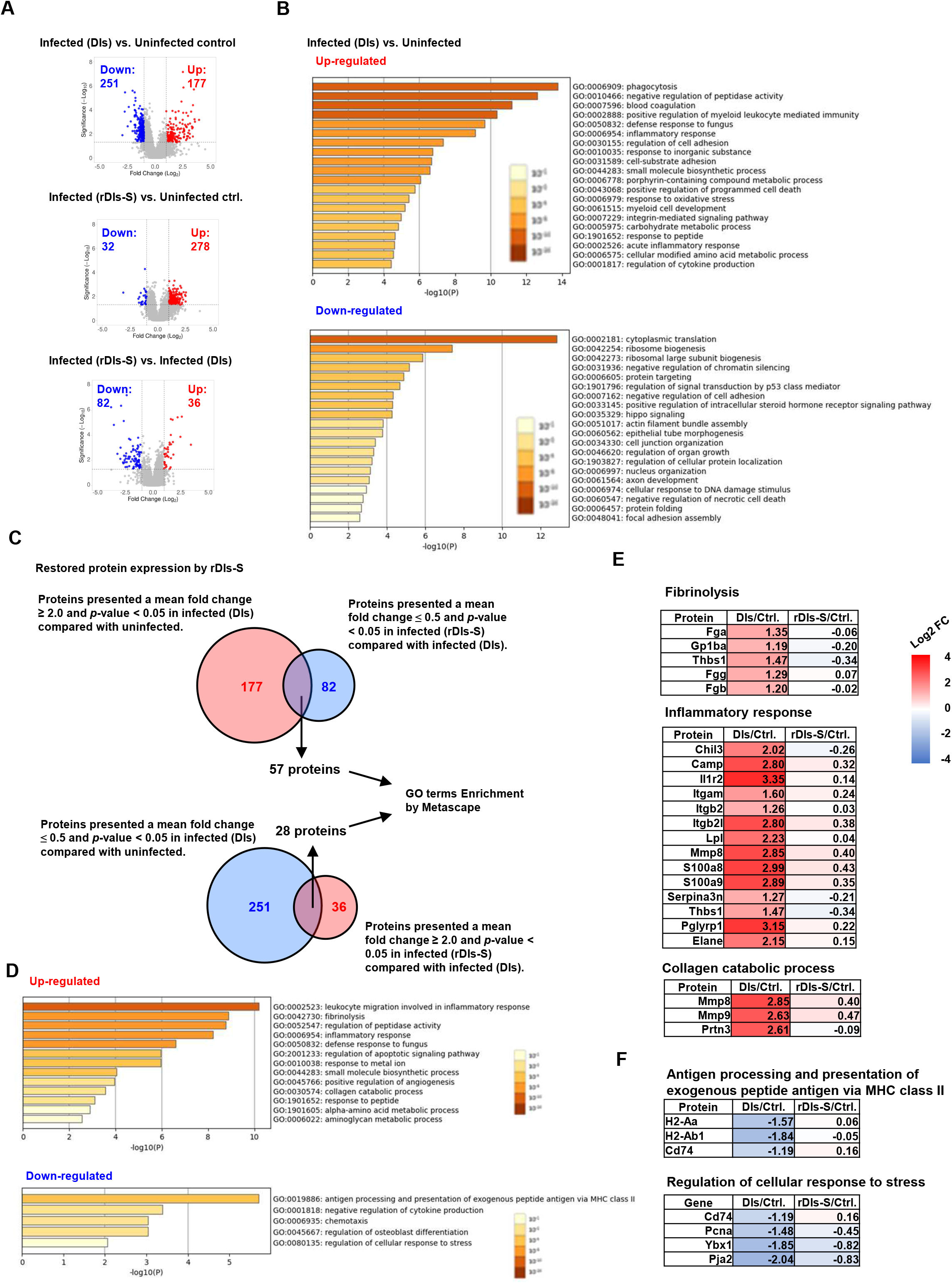
Proteomic analysis in vaccinated mice after SARS-CoV-2 infection. The protein expression levels in the lung tissues of hACE2 transgenic mice 7 days after challenge infection were analyzed using multiplex peptide labeling and MS. (A) Volcano plot for differentially expressed proteins. Comparison of the protein expression levels between uninfected mice and infected mice inoculated with DIs (top), between uninfected mice and infected mice inoculated with rDIs-S (middle), and between infected mice inoculated with DIs and infected mice inoculated with rDIs-S (bottom). x-axis: expression ratios. y-axis: *p-*values of the comparisons. Vertical dotted lines indicate a 2-fold increase or decrease in the protein expression level in mice inoculated with DIs or rDIs-S. Horizontal lines indicate a *p*-value of 0.05 from a chi-squared test. Red circles: proteins with concentration increases of more than 2.0-fold in each comparison (DIs vs. Uninfected, rDIs-S vs. Uninfected, and rDIs-S vs. DIs); blue circles: proteins with concentrations decreased to less than half in each comparison (DIs vs. Uninfected, rDIs-S vs. Uninfected, and rDIs-S vs. DIs). (B) The top-20 gene ontology (GO) enrichment terms related to biological processes (BP) of the proteins that were upregulated (upper) and downregulated (lower), as analyzed by Metascape. (C) Number of proteins with altered expression in infected mice without vaccination for which expression was restored by rDIs-S vaccination. (D) GO enrichment terms related to BP of the genes encoding the proteins that were restored by rDIs-S vaccination among the upregulated (upper) and downregulated (lower) proteins in the infected (DIs) mice. (E) Representative cluster of GO enrichment terms related to BP of the genes encoding the proteins that were restored by rDIs-S vaccination, among the upregulated proteins in the infected (DIs) mice. (F) Representative cluster of GO enrichment terms related to BP of the genes encoding the proteins that were restored by rDIs-S vaccination, among the downregulated proteins in the infected (DIs) mice.

### Long-term humoral immune responses following vaccination with rDIs-S

To investigate the ability of rDIs-S to establish a long-lived immunological memory, 8-week-old BALB/c mice were vaccinated twice at a 3-week interval, and the antibody responses specific to S protein were monitored by enzyme-linked immunosorbent assay (ELISA) using the S protein ectodomain trimer as an antigen. As shown in Fig. 6, S protein-specific IgG was detected 3 weeks after the 1^st^ vaccination, and the IgG titer was increased significantly after the 2^nd^ vaccination. Importantly, the titer of S protein-specific IgG was maintained at the same level from 4 to 24 weeks after the 2^nd^ vaccination (7 to 27 weeks after the 1^st^ vaccination), indicating that the titers of S protein-specific IgG induced by rDIs-S were maintained for at least 6 months after vaccination. This result raises the possibility that rDIs-S confers long-term protection against SARS-CoV-2 infection ^23^.

**Fig. 6.**
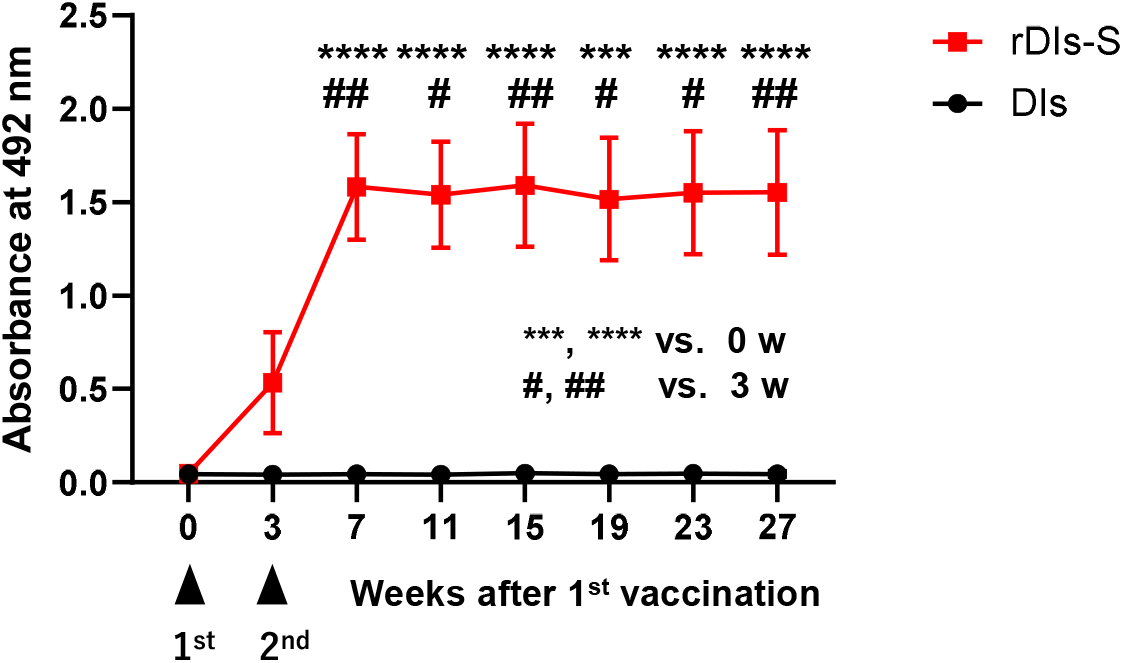
Time course of antibody responses after vaccination with rDIs-S in mice. BALB/c mice were inoculated twice intradermally with 1 × 10^8^ PFU of rDIs-S or DIs at 3-week intervals. Time course of the production of IgG specific for SARS-CoV-2 S protein as measured by ELISA (n =12 per group). *p*-values were calculated using a two-tailed Kruskal-Willis test, followed by a Dunn’s multiple comparison test.

## Discussion

This work demonstrated the efficacy of rDIs-S, an attenuated vaccinia virus vaccine engineered to encode the SARS-CoV-2 S protein, against SARS-CoV-2 infection; the efficacy was assessed in mouse and macaque models. Two vaccinations with rDIs-S induced nAbs against not only the “classic” (original) SARS-CoV-2 strain isolated in early 2020 but also variant strains, while also inducing IFN-γ-producing T cells specific for SARS-CoV-2 S antigen. These effects resulted in a reduction of SARS-CoV-2 virus titers, along with protection from lethal infection in hACE2 transgenic mice and protection from pneumonia in cynomolgus macaques. A comprehensive analysis of protein levels in SARS-CoV-2-infected infected mice showed that the expression of proteins involved in tissue damage and inflammation was attenuated in the DIs-S-vaccinated mice compared to the DIs-immunized animals.

In the present study, we immunized hACE2 transgenic mice and the cynomolgus macaques twice with rDIs-S. After the 2^nd^ vaccination, nAb titers against SARS-CoV-2 and the amounts of IFN-γ produced by T lymphocytes were increased from the pre-immunization baseline levels. These observations suggest that rDIs-S will be immunogenic in people who previously have been immunized with the attenuated vaccinia virus ^24^, and a booster effect from repeated vaccination is expected to enhance and maintain immunological memory against SARS-CoV-2. This effect may make rDIs-S advantageous compared to SARS-CoV-2 vaccines employing other virus vectors, such as the adenovirus vaccine encoding the SARS-CoV-2 S protein that is recommended as a single-dose vaccination. We note, however, that a booster effect was reported in aged mice vaccinated twice with the adenovirus vaccine carrying the SARS-CoV-2 *S* gene ^25,26^.

The results of the present study, including the induction of nAbs against the early-pandemic SARS-CoV-2 strain and protective efficacy, are consistent with the results of previous studies in which mice ^27–31^ and rhesus macaques ^32^ were immunized with modified vaccinia Ankara strains carrying the SARS-CoV-2 *S* gene. Those studies, like ours, confirmed the safety of vaccinia-based vaccines and their immunogenicity in animals vaccinated repeatedly, indicating that vaccinia-based vaccines may be usable even in younger populations and in the elderly with preexisting immunity against smallpox ^24^. Furthermore, our results demonstrated the efficacy of rDIs-S against variant strains, since neutralization activity was seen against the variant strains in macaques, and an improved survival rate was seen in hACE2 transgenic mice, a model that had not been examined in other studies. Therefore, we expect that rDIs-S will confer broad protection against multiple variants of SARS-CoV-2 ^33^.

Using TMT-based quantitative proteomic analysis of lung homogenates from uninfected, DIs-immunized, and rDIs-S-immunized mice, we found that inoculation with rDIs-S protected the mice from the severe pathogenic effects of SARS-CoV-2 infection, such as tissue destruction, inflammation, coagulation, fibrosis, and angiogenesis. These changes in protein expression, which were observed in the control (DIs-immunized) mice after infection with SARS-CoV-2, also are seen in critical COVID-19 patients ^34^, indicating the utility of the hACE2 transgenic mouse model for evaluating the potential protective efficacy of vaccines against severe COVID-19 symptoms. In addition, since the TMT-proteomic analysis detects changes in protein levels in a comprehensive and sensitive manner, this technology also may serve as a safety evaluation system to identify factors related to the adverse events that have been seen with the current vaccines ^35^. Of note, coagulation factors are thought to be activated by COVID-19 and vaccination ^35,36^, but the expression levels of the coagulation factors in the rDIs-S-vaccinated mice were comparable to those in the uninfected mice (Fig. 5E).

We demonstrated that rDIs-S provides long-lived humoral immune responses for at least 6 months after vaccination in mice (Fig. 6). Recent reports have shown that the antibody levels induced by the current mRNA vaccine decline dramatically at 6 months after the 2^nd^ vaccination ^23,37^. Thus, additional periodic vaccination would be required for the control of COVID-19 using the current vaccines. However, since the current mRNA vaccines may cause undesirable adverse events, the long-term immune memory response conferred by rDIs-S may be one of the useful advantages for the development of new vaccines.

In the present study, we demonstrated the efficacy of rDIs-S, an attenuated vaccinia virus carrying the SARS-CoV-2 *S* gene. Furthermore, given that vaccination with rDIs-S effectively induced antibody and T-lymphocyte responses that also reacted with variant strains, rDIs-S may be useful for conferring protection against new variants by use as a booster after vaccination with 1^st^-generation SARS-CoV-2 vaccines.

## Methods

### Ethics statement

This study was carried out in strict accordance with the “Guidelines for the Husbandry and Management of Laboratory Animals” of the Tokyo Metropolitan Institute of Medical Science and the Research Center for Animal Life Science at Shiga University of Medical Science, and with the “Standards Relating to the Care and Fundamental Guidelines for Proper Conduct of Animal Experiments and Related Activities in Academic Research Institutions” under the jurisdiction of the Ministry of Education, Culture, Sports, Science and Technology of Japan. The animal experimental protocols were approved by the Ethics Committee of Animal Experiments of the Toyo Metropolitan Institute of Medical Science (Permission Nos. 20-85, 20-86, 21-79, and 21-080), and by the Shiga University of Medical Science Animal Experiment Committee (Permit No. 2020-6-20). In the macaque study, regular veterinary care and monitoring, balanced nutrition, and environmental enrichment were provided by personnel of the Research Center for Animal Life Science at Shiga University of Medical Science. The macaques were euthanized at the endpoint (7 days after SARS-CoV-2 virus inoculation) using ketamine/xylazine followed by intravenous injection of pentobarbital (200 mg/kg). Animals were monitored every day during the study to permit calculation of clinical scores (Supplemental Table 3) and underwent daily veterinary examinations intended to help alleviate suffering. The animals were euthanized if their clinical score reached 15 (a humane endpoint).

### Cells

Primary chicken embryonic fibroblasts were prepared for constructing and propagating the recombinant VACV DIs strain that carries the gene encoding the spike protein of SARS-CoV-2. Seven-day-old chicken embryos were collected in Hanks’ Balanced Salt Solution [HBSS (-)] supplemented with 50 U/mL penicillin, 50 μg/mL streptomycin, and 0.1% glucose. After removing the eyes, brain, beak, wings, and feet from each embryo, the rest of the body was minced with scissors and digested in TrypLE Select (Thermo Fisher Scientific, Waltham, MA, USA). The resulting chicken embryonic fibroblasts were grown in Dulbecco’s Modified Eagle Medium (DMEM; Nacalai Tesque, Kyoto, Japan) supplemented with 10% fetal calf serum and tryptose phosphate broth.

### Vaccine construction

Codon optimization was performed for the spike protein-encoding gene sequence of SARS-CoV-2 (AI/I-004/2020 strain GISAID EPI_ISL_407084, or Delta variant hCoV/Japan/TY11-927-P1/2021 strain GISAID EPI_ISL_2158617) to facilitate stable expression in the context of DIs. Silent mutations were introduced in the sequences encoding nCoV-S to remove stop signal sequences (TTTTTNT) for the vaccinia virus early promoter. The resulting synthetic DNA encodes a modified nCoV-S (mnCoV-S) or Delta variant-S and was designed to include flanking SbfI and AsiSI restriction sites upstream and downstream (respectively) of the *S* open reading frame (ORF). This synthetic DNA was cloned into the DIs vector plasmid pSMART-DIs-L3-GPTF (purchased from GenScript, Nanjing, China), which harbors the *Escherichia coli gpt* gene (encoding xanthine-guanine phosphoribosyltransferase, XGPRT) under control of the VACV *p7.5* early promoter. The resulting constructs, designated pSMART-DIs-L3-mnCoV-S-GPTF and pSMART-DIs-L3-mnCoV-Delta S-GPTF, were linearized by digestion with the ApaI restriction enzyme. The linearized plasmid was purified and transfected into primary chicken embryonic fibroblasts that had been infected with DIs at a multiplicity of infection of 10 for 1 h. After 20 h, the virus-cell mixture was harvested by scraping of the cell layer, and the resulting suspension was frozen at −80 °C until use. rDIs-mnCoV-S and rDIs-mnCoV-Delta S (i.e., the rDIs-S and rDIs-S (Delta) used in this study) were purified in the presence of the selective reagent mycophenolic acid, an inhibitor of purine metabolism; the use of a vector containing the *E. coli gpt* gene in the presence of xanthine and hypoxanthine permitted the cultures to overcome the blocking of the pathway for GMP synthesis caused by mycophenolic acid, as described previously ^38^. DIs was used as a control virus.

### SARS-CoV-2

SARS-CoV-2 JP/TY/WK-521/2020 (WK-521) (GenBank Sequence Accession: LC522975), SARS-CoV-2 hCoV-19/Japan/TY8-612/2021 (TY8-612) (GISAID strain name: EPI_ISL_1123289), SARS-CoV-2 hCoV/Japan/TY11-927-P1/2021 (TY11-927) (GISAID strain name: EPI_ISL_2158617), and SARS-CoV-2 hCoV-19/Japan/TY38-873/2021 (TY38-873) (GISAID strain name: EPI_ISL_7418017) were used as challenge strains; these isolates were kindly provided by Drs. Masayuki Saijo, Mutsuyo Takayama-Ito, Masaaki Sato, and Ken Maeda, National Institute of Infectious Disease (NIID) ^39^. SARS-CoV-2 JPN/Shiga/SUMS2/2020 (SUMS2) (GISAID strain name: EPI_ISL_10434280), which was isolated from a patient hospitalized in the Shiga University of Medical Science Hospital, encodes an S protein with a D614G substitution. All virus strains used in this study are listed in Supplemental Fig. 4. The nucleotide sequence of WK-521 has 99.9% similarity to that of the Wuhan-Hu-1 strain (NC_045512.2). The WK-521 virus was propagated twice at the NIID, and then once at the Shiga University of Medical Science or twice at The Tokyo Metropolitan Institute of Medical Science, using the VeroE6/TMPRSS2 cell line (JCRB Cell Bank, Osaka, Japan). The other variants also were propagated at the NIID, and then once at the Tokyo Metropolitan Institute of Medical Science or at the Shiga University of Medical Science, using VeroE6/TMPRSS2. The macaques were challenged with the WK-521 virus (2 × 10^7^ mean tissue culture infectious dose (TCID_50_)), which was inoculated into the animals’ conjunctiva (0.05 mL × 2), nostrils (0.5 mL × 2), oral cavity (0.9 mL), and trachea (5 mL) with pipettes and catheters; the animals were placed under ketamine/xylazine anesthesia prior to inoculation. Experiments using the virus were performed in the Biosafety Level 3 facility of the Research Center for Animal Life Science, the Shiga University of Medical Science, and the Tokyo Metropolitan Institute of Medical Science.

VeroE6/TMPRSS2 cells were grown in DMEM supplemented with 10% inactivated fetal bovine serum (Capricorn Scientific GmbH, Ebsdorfergrund, Germany), penicillin (100 units/mL), streptomycin (100 μg/mL), and G418 (1 mg/mL; Nacalai Tesque). To assess viral replication, serial dilutions of swab samples and tissue homogenate samples (10% w/v in HBSS) were inoculated onto confluent VeroE6/TMPRSS2 cells. The VeroE6/TMPRSS2 cells were cultured in DMEM supplemented with 0.1% bovine serum albumin (BSA), penicillin, streptomycin, and gentamycin (50 μg/mL; Fujifilm, Tokyo, Japan). Cytopathic effects were determined by examination under a microscope 3 days later.

### Mice

C57BL/6J mice were purchased from CLEA Japan, Inc. (Tokyo, Japan). hACE2 transgenic mice were obtained from the National Institute of Biomedical Innovation, Health and Nutrition as ACE2 Tg #17 (Strain nbio0298). To maintain the heterozygouse (hACE2 Tg/+) hACE2 mice, C57BL/6 mice and heterozygouse (hACE2 Tg/+) hACE2 mice were mated. BALB/c mice were purchased from Japan SLC, Inc. (Hamamatsu, Japan). Throughout the mouse studies, animals were provided with free access to food and water, and were maintained on a 12-h light/12-h dark cycle. Prior to inoculation, animals were anesthetized by intraperitoneal administration of a ketamine/xylazine mixture. Animals then were inoculated intratracheally with 20 TCID_50_ per 50 μL of the WK-521 strain, 100 plaque-forming units (PFU) per 50 μL of the TY8-612 strain, or 50 PFU per 50 μL of the TY8-612 strain. We recently generated an adenoviral vector expressing the hACE2-encoding gene under the *EF1α* promoter with a leftward orientation (rAd5 pEF1α-hACE2-L) as a novel transgene vector to confer SARS-CoV-2 susceptibility in wild-type mice ^20^. Using this model, we investigated the ability of rDIs-S (encoding spike derived either from an early-pandemic strain of SARS-CoV-2 or from a Delta variant) to provide protection against an Omicron variant. BALB/c mice were inoculated intranasally with 5 × 10^7^ focus-forming units (FFU) per 50 μL of rAd5 pEF1-hACE2-L. Five days after the inoculation, the BALB/c mice were inoculated intranasally with 1 × 10^5^ PFU per 50 μL of the TY38-837 strain of SARS-CoV-2. Body weight was monitored daily; mice that lost 30% or more of their initial body weight were humanely euthanized and scored as dead.

### Macaques

Nine- to 18-year-old female and male cynomolgus macaques that were born in the Philippines or at the Shiga University of Medical Science were used; for animals bred in-house, the maternal macaques originated from Vietnam, and the paternal macaques originated from Indonesia or China. All procedures were performed under ketamine/xylazine anesthesia, and all efforts were made to minimize suffering. Food pellets of CMK-2 (CLEA Japan, Inc.) were provided once per day after recovery from the anesthesia, and drinking water was available *ad libitum.* The animals were single-housed in cages equipped with climbable bars for environmental enrichment under controlled conditions of humidity (46% to 70%), temperature (22.3 to 23.9 °C), and light (12-h light/12-h dark cycle, lights on at 8:00 a.m.). Two weeks before virus inoculation, two temperature data loggers (iButton, Maxim Integrated, San Jose, CA, USA) were implanted in the peritoneal cavity of each macaque under ketamine/xylazine anesthesia followed by isoflurane inhalation; the data loggers permitted monitoring of body temperature. The macaques used in the present study were confirmed, by testing, to be free of herpes B virus, hepatitis E virus, *Mycobacterium tuberculosis, Shigella* spp., *Salmonella* spp., and *Entamoeba histolytica.* Attenuated VACV (1 × 10^8^ PFU) was injected intracutaneously twice using a syringe with a 29-G needle. Macaques were distinguished by identification numbers as follows: C1-C4, macaques inoculated with DIs; V1-V4, macaques inoculated with rDIs-S.

Using animals anesthetized with ketamine/xylazine, two cotton sticks (Eiken Chemical, Ltd., Tokyo, Japan) were used to collect fluid samples from the conjunctiva, nasal cavity, oral cavity, trachea and rectum, and the sticks were subsequently immersed in 1 mL of DMEM supplemented with 0.1% BSA and antibiotics. A bronchoscope (MEV-2560; Machida Endoscope Co., Ltd., Tokyo, Japan) and cytology brushes (BC-203D-2006; Olympus Co., Tokyo, Japan) were used to obtain bronchial samples. Samples were collected on the indicated days.

Chest X-ray radiographs were obtained using the I-PACS system (Konica Minolta Inc., Tokyo, Japan) and a PX-20BT mini (Kenko Tokina Corporation, Tokyo, Japan). Saturation of peripheral oxygen (SpO2) was measured with a pulse oximeter (Nellcor^™^; Medtronic plc, Dublin, Ireland).

### Extraction of RNA and <u>q</u>uantitative reverse transcription polymerase chain reaction (qRT-PCR) for the detection of SARS-CoV-2 RNA

Total RNA samples were extracted from swab samples and tissue samples from the macaques using the QIAamp Viral RNA Mini Kit and RNeasy Plus Mini Kit (Qiagen, Venlo, Netherlands) according to the manufacturer’s instructions. The levels of RNA corresponding to the N protein-encoding gene of SARS-CoV-2 were measured using the TaqMan Fast Virus 1-step Master Mix (Thermo Scientific). Each 20-μL reaction mixture contained 5.0 μL of 4× TaqMan Fast Virus 1-Step Master Mix, 0.25 μL of 10 μM probe, 1.0 μL each of 10 μM forward and reverse primers, 7.75 μL of nuclease-free water, and 5.0 μL of nucleic acid extract. Amplification was carried out in 96-well plates using a CFX-96 cycler equipped with CFX Maestro software (Bio-Rad, Hercules, CA, USA). The thermocycling conditions were as follows: 5 min at 50 °C for reverse transcription, 20 s at 95 °C for the inactivation of reverse transcriptase and initial denaturation, and 45 cycles of 5 s at 95 °C and 30 s at 60 °C for amplification. Each run included a no-template control reaction as well as reactions intended to provide a standard curve. The latter used *in vitro* transcribed RNA of the N protein-encoding gene (at 10^0^, 10^1^, 10^2^, 10^3^, 10^4^, 10^6^, and 10^8^ copies/reaction); this template was generated from the cDNA of SARS-CoV-2 AI/I-004/2020 using the T7 RiboMAX Express Large Scale RNA Production System (Promega, Madison, WI, USA). The primers and probe used in this study were as follows: forward primer, 5’-GACCCCAAAATCAGCGAAAT-3’; reverse primer, 5’-TCTGGTTACTGCCAGTTGAATCTG-3’; and probe, 5’-(FAM)-ACCCCGCATTACGTTTGGTGGACC-(BHQ-1)-3’, where FAM and BHQ-1 are 6-fluorescein amidite and Black Hole Quencher-1, respectively.

### Histopathological examination

Lungs were obtained at necropsy, and eight lung tissue slices were collected from each macaque: one slice from each upper lobe and middle lobe, and two slices from each lower lobe in the bilateral lungs. These slices were fixed in 10% neutral buffered formalin for approximately 72 h, embedded in paraffin, and cut into 3-μm-thick sections, which were mounted on glass slides. Sections were stained with hematoxylin and eosin and observed by light microscopy. Histological evaluation was performed by two pathologists, both blinded to sample identification, based on criteria established for influenza virus infection ^40^ as follows: 0, normal lung; 1, mild destruction of bronchial epithelium; 2, mild peribronchiolar inflammation; 3, inflammation in the alveolar walls resulting in alveolar thickening; 4, mild alveolar injury accompanied by vascular injury; 5, moderate alveolar injury and vascular injury; and 6 and 7, severe alveolar injury with hyaline membrane-associated alveolar hemorrhage (under or over 50% of the section area, respectively). The mean score for the eight sections was calculated for each macaque, and the mean score of the two pathologists was defined as the histological score. After autoclaving the slides in citrate buffer (pH 9) for antigen retrieval, SARS-CoV-2 N antigen was detected using monoclonal anti-N antibody 8G8A (Bioss, Inc., Boston, MA, USA) and a secondary antibody.

### Blood cytokine and biochemical analyses

Levels of cytokines/chemokines in macaque plasma were measured using the Milliplex MAP Non-human Primate Cytokine Panel in combination with a Luminex 200 (Millipore Corp., Billerica, MA, USA) according to the manufacturer’s instructions.

### Virus neutralization assay

Complement in plasma samples was inactivated by heating at 56 °C for 1 h. The diluted samples were mixed for 30 min with 100 TCID50/well of the SARS-CoV-2 strains shown in Supplemental Fig. 4. Then, each mixture was added onto a VeroE6/TMPRSS2 monolayer in 96-well plates. After 1 h of incubation, the cells were cultured in DMEM supplemented with 0.1% BSA. After incubation at 37 °C for 3 days, the number of wells showing a cytopathic effect (CPE) was counted. Neutralization titers are expressed as the dilution at which CPEs were observed in 50% of the wells. This assay was performed in quadruplicate culture.

### Detection of cytokine-producing cells by enzyme-linked immunosorbent spot (ELISpot)

After separation from red blood cells, peripheral blood mononuclear cells (PBMCs) were stored at −80 °C until use. Thawed cells (5 × 10^5^/well) were cultured overnight with a peptide pool of SARS-CoV-2 S protein (0.6 nmol/mL; PepTivator; Miltenyi Biotech, Bergisch Gladbach, Germany) in the presence of anti-CD28 antibody (0.1 μg/mL); culturing was performed in ELISpot plates coated with anti-IFN-γ and anti-IL-2 antibodies (Cellular Technology, Ltd., Shaker Heights, OH, USA). The number of cytokine-producing cells was counted according to the manufacturer’s instructions. Stimulation indices (S.I.) were calculated as follows: S.I. = number of spots in the culture of cells with peptides / number of spots in the culture of cells without peptides. In the mouse study, isolated single splenocytes were used for the ELISpot assay. The splenocytes were cultured with 1 μg/mL of the peptide pool of SARS-CoV-2 S protein (PepTivator SARS-CoV-2 Prot_S, SARS-CoV-2 Prot_S1, and Prot_S+, which cover the full-length of the S protein; Miltenyi Biotech) for 24 h in ELISpot plates coated with anti-mouse-IFN antibody (Mabtech AB, Nacka Strand, Sweden). The procedures were carried out according to the manufacturer’s instructions (Mabtech, #3321-2H). After drying the ELISpot plates, the number of spots in each well was counted with an automated ELISpot plate reader (Advanced Imaging Devices GmbH, Strassberg, Germany).

### *In vivo* cytotoxic T lymphocyte (CTL) assay

Single cells were isolated from the spleens of C57BL/6J mice. After the lysis of red blood cells, splenocytes were incubated with 1 μg/mL of the peptide pool of SARS-CoV-2 S protein (PepTivator, as above) in a 37 °C water bath for 1 h. After washing with HBSS, unpulsed splenocytes and the S-peptide-pulsed splenocytes were stained with 0.5 μM and 5 μM (respectively) of 5-(and 6-) carboxyfluorescein diacetate succinimidyl ester (CFSE; BioLegend, San Diego, CA, USA). The unpulsed and peptide-pulsed splenocytes were mixed at a 1:1 ratio (5 × 10^6^ cells each), and the mixture was injected intravenously into DIs- and rDIs-S-inoculated mice. Twenty-four hours later, the spleens were harvested, and the percentages of cells positive for staining with CFSE (i.e., CFSE^+^ cells) that were CFSE^low^ and CFSE^hi^ were assessed by flow cytometry. The percent specific killing was calculated as: 1 – (Non-transferred control ratio / Experimental ratio)] × 100.

### Detection of IgG specific for the SARS-CoV-2 spike protein

For enzyme-linked immunosorbent assay (ELISA), recombinant SARS-CoV-2 spike S1+S2 extracellular domain (ECD) (Sino Biological, Inc., Beijing, China) was coated onto 96-well round-bottom plates, and the plates were incubated overnight at 4 °C. The plates were blocked with 1% BSA in phosphate-buffered saline [PBS (-)] containing 0.5% Tween 20 and 2.5 mM ethylenediaminetetraacetic acid, then incubated with a 500-fold dilution of sera from C57BL/6 and hACE2 transgenic mice immunized with either rDIs-S or DIs, a 1000-fold dilution of sera from BALB/c mice immunized with either rDIs-S or DIs, or a 1000-fold dilution of plasmas from cynomolgus macaques immunized with either rDIs-S or DIs. After extensive washing, the plates were incubated with horseradish peroxidase-conjugated sheep anti-mouse IgG polyclonal antibody (GE Healthcare, Chicago, IL, USA). Antigen-antibody interactions were detected using *o*-phenylenediamine solution as the substrate (Nacalai Tesque), and the binding activity was measured by monitoring absorbance at 490 nm. For the bead array assay to detect IgG specific for S1, RBD, and S2 in plasma, the Milliplex SARS-CoV-2 Antigen Panel 1 IgG was used according to the manufacturer’s instructions (Merck Millipore, Darmstadt, Germany).

### TMTpro llplex MS analysis

Lysates extracted from mouse lung tissues with a bead shocker were processed and digested using an EasyPep Mini MS Sample Prep Kit (Thermo Fisher Scientific) according to the manufacturer’s protocol. Then, 25 μg of peptides from each sample were labeled with 0.25 mg of the TMTpro TMT-labeling reagent (Thermo Fisher Scientific) according to manufacturer’s protocol. After TMT labeling, aliquots from the 11 sample channels were combined in an equal ratio, dried using a vacuum concentrator, and resuspended in 0.1% trifluoroacetic acid (TFA). Samples were separated into eight fractions using a High-pH Reversed-Phase Peptide Fractionation Kit (Thermo Fisher Scientific) according to the manufacturer’s protocol. Then, 1 μg of peptides from each fraction were analyzed by liquid chromatography coupled to tandem mass spectrometry (LC-MS/MS) on an EASY-nLC 1200-connected Orbitrap Fusion Lumos Tribrid MS (Thermo Fisher Scientific) equipped with a High-Field Asymmetric Waveform Ion Mobility Spectrometry (FAIMS) -Pro ion mobility interface (Thermo Fisher Scientific). Peptides were separated on an analytical column (C18, 1.6-μm particle size × 75 μm diameter × 250 mm; Ion Opticks, VIC, Australia) using a gradient of 0% to 28% acetonitrile over 240 min at a constant flow rate of 300 nL/min. Peptide ionization was performed using a Nanospray Flex Ion Source (Thermo Fisher Scientific). The FAIMS-Pro was set to three phases (−40, −60, and −80 CV); a ‘ 1-s cycle for a phase’ data-dependent acquisition method was used, in which the most intense ions for every 1-s interval were selected for MS/MS fragmentation by higher-energy C-trap dissociation (HCD). MS raw files were analyzed using the Sequest HT search program in Proteome Discoverer 2.4 (Thermo Fisher Scientific). MS/MS spectra were searched against the Swiss-Prot reviewed mouse reference proteome database (UniProt). TMTpro-based protein quantification was performed using the Reporter Ions Quantifier node in Proteome Discoverer 2.4.

### Volcano plot

The volcano plot was prepared using VolcaNoseR software ^41^.

### Enrichment analysis

GO term enrichment related to BP was analyzed by Metascape (http://metascape.org) ^22^. Terms with a *p*-value <0.05, a minimum count of 3, and an enrichment factor >1.5 were collected and grouped into clusters based on their membership similarities.

### Statistical analyses

Data plotted on a linear scale are expressed as the mean ± standard deviation (SD), except for the mean ± standard error of mean (SEM) of body weight change in Fig. 1J. Data plotted on logarithmic scales are expressed as the geometric mean ± geometric SD. Inferential statistical analysis was performed using a two-tailed non-paired Student’s *t*-test, Mann-Whitney U test, One-way ANOVA followed by Tukey’s test, or chi-squared test, as appropriate. Statistical significance was set at *p* < 0.05. The Prism software package (version 9.1; GaphPad Software, San Diego, CA, USA) was used for all statistical analyses.

## Acknowledgments

This work was supported, in part, by funding from the Japan Agency for Medical Research and Development (AMED) under Grant Nos. 19fk0108172, 20nk0101615, 20fk0108410, and 20fk0108538; and by a grant from the Tokyo Metropolitan Government. Misako Nakayama was supported by the Naito Foundation. Cong Thanh Nguyen was supported by the Sato Yo International Scholarship Foundation. We thank Drs. Masayuki Saijo, Mutsuyo Takayama-Ito, Masaaki Sato, and Ken Maeda for providing the SARS-CoV-2 strains; Drs. Hideaki Tsuchiya, Ikuo Kawamoto, Takahiro Nakagawa, and Iori Itagaki for animal care; and Hideaki Ishida, Naoko Kitagawa, Takako Sasamura, Chikako Kinoshita, and Sayaka Ono for assistance in the experiments.

## Author contributions

Y.I. and M.K. conceived the study. H.I., F.Y., M.N., A.E., N.Y., K.Y., C.T.N., Y.K., T.H., T.S., M.H., S.T., A.T., Y.M., K.H., and M.S. performed the experiments. K.I. and Y.S. supervised the study. H.I., F.Y., M.N., A.E., N.Y., C.T.N., T.H., T.S., Y.M., Y.S., Y.I., and M.K. participated in data analysis, interpretation, and manuscript review. Y.I., F.Y., and M.K. wrote the manuscript.

## Competing interest statement

The authors declare no competing interests.

## Notes

### Competing Interest Statement

The authors have declared no competing interest.

