## Supplemental Tables and figures for "An attenuated vaccinia vaccine encoding the SARS-CoV-2 spike protein elicits broad and durable immune responses, and protects cynomolgus macaques and human ACE2 transgenic mice from SARS-CoV-2 and its variants"

Supplemental Table 1

**Supplemental Table 1. Infectious virus titers in swab samples.**

| Virus titers ($log_{10}$ $TCID_{50}$/mL) | | | | | | |
|---|---|---|---|---|---|---|
| Sample | Vaccine | Animal | Days after virus inoculation [1] | | | |
| | | | 1 | 3 | 5 | 7 |
| Nasal Cavity | DIs | C1 | 3.50 | 2.50 | < [2] | 1.50 |
| | | C2 | 3.77 | < 1.33 [3] | < | < 1.50 [4] |
| | | C3 | 4.00 | < | < | < |
| | | C4 | 3.67 | < 0.67 [5] | < 0.67 | < |
| | rDIs-S | V1 | 3.00 | < | < | < |
| | | V2 | < | < | < | < |
| | | V3 | 2.67 | < | < | < |
| | | V4 | 4.50 | < | < | < |
| Oral cavity | DIs | C1 | 1.67 | < 0.67 | < 1.23 [6] | < |
| | | C2 | < | < | < | < |
| | | C3 | < 0.83 [7] | < | < | < |
| | | C4 | < | < | < | < |
| | rDIs-S | V1 | < | < | < | < |
| | | V2 | 2.50 | < | < | < |
| | | V3 | < | < | < | < |
| | | V4 | < | < | < | < |
| Trachea | DIs | C1 | 4.50 | 1.50 | < 1.00 [8] | < 1.50 |
| | | C2 | 3.50 | < | < | < |
| | | C3 | 3.33 | < | < | < |
| | | C4 | 3.33 | < | < | < |
| | rDIs-S | V1 | < | < | < | < |
| | | V2 | < | < | < | < |
| | | V3 | < 1.00 | < | < | < |
| | | V4 | 2.33 | < | < | < |
| Bronchus | DIs | C1 | < 0.67 | < | < | < |
| | | C2 | 3.33 | < | < | < |
| | | C3 | 3.67 | < | < | < |
| | | C4 | 2.33 | < | < | < |
| | rDIs-S | V1 | < | < | < | < |
| | | V2 | < | < | < | < |
| | | V3 | < | < | < | < |
| | | V4 | < 0.67 | < | < | < |

[1] Swab samples were collected on the indicated days.

[2] <: Virus titers were below the limit of detection (0.67 $log_{10}TCID_{50}$/mL).

[3] < 1.33: Three CPE-positive wells were detected in quadruplicate cultures of undiluted samples.

[4] < 1.50: Three CPE-positive wells were detected in quadruplicate cultures of undiluted samples.

[5] < 0.67: One CPE-positive well was detected in quadruplicate cultures of undiluted samples.

[6] < 1.23: Two and one CPE-positive wells were detected in quadruplicate cultures of undiluted samples and 1/10 diluted samples, respectively.

[7] < 0.83: One CPE-positive well each was detected in quadruplicate cultures of undiluted samples and 1/10 diluted samples.

[8] < 1.00: Two CPE-positive wells were detected in quadruplicate cultures of undiluted samples.

Supplemental Table 2

**Supplemental Table 2. Infectious virus titers in lung tissue samples.**

| Virus titers ($\log_{10}$ $TCID_{50}$/0.1 g tissue[1]) | | | | | | | |
|---|---|---|---|---|---|---|---|
| Vaccine | Animal | right upper | right middle | right lower | left upper | left middle | left lower |
| DIs | C1 | < 0.67 [2] | < 1.33 [3] | < 2.00 [4] | < [5] | < | < |
| | C2 | < | < | < 0.67 | < | < 1.33 | < |
| | C3 | < | < 0.83 [6] | < | < | < 0.67 | < |
| | C4 | < | < | < | < | < | < |
| rDIs-S | V1 | < | < | < | < | < | < |
| | V2 | < | < | < | < | < | < |
| | V3 | < | < | < | < | < | < |
| | V4 | < | < | < | < | < | < |

[1] Lung tissues were collected following euthanasia at 7 days after virus inoculation. The tissues were homogenized to prepare a 10% w/v solution.

[2] < 0.67: One CPE-positive well was detected in quadruplicate cultures of undiluted samples.

[3] < 1.33: Three CPE-positive wells were detected in quadruplicate cultures of undiluted samples.

[4] < 2.00: Two CPE-positive wells were detected in quadruplicate cultures of undiluted samples.

[5] <: Virus titers were below the limit of detection (0.67 $\log_{10}TCID_{50}$/0.1 g tissue).

[6] < 0.83: One CPE-positive well each was detected in quadruplicate cultures of undiluted samples and 1/10 diluted samples.

Supplemental Table 3

**Supplemental Table 3. Clinical scoring used in the present study**

| Parameter | Degree of parameter | Possible score |
|---|---|---|
| Fever | Normal (< 39 °C) | 0 |
| | Elevated temperature (39-40 °C) | 3 |
| | High temperature (> 40 °C) | 5 |
| Posture | Piloerection of body hair | 1 |
| | Decreased activity, decreased normal behavior/Occasionally lying down, huddled, active when people in room | 2 |
| | Huddled on camera, active when people in room/Lying down, getting up when approached, using cage for support | 3 |
| | Huddled when people in room, shaking, toes and hands clenched/Lying down, not getting up when approached or prompted | 5 |
| Respiration | Increased or decreased; mild cough and clear nasal discharge | 3 |
| | Labored breathing through mouth; severe cough and severe nasal discharge | 5 |
| Appetite | Slightly decreased | 1 |
| | Decreased | 2 |
| | Severely decreased | 5 |
| Skin | Flushed appearance | 2 |
| | Visible rash | 2 |
| | Bleeding | 5 |

Animals were monitored every day during the study to be clinically scored.
Animals were euthanized if their clinical scores reached 15 (a humane endpoint).

Supplemental Table 4

**Supplemental Table 4. SARS-CoV-2 used in this study**

| Abbr. | Strain name | GISAID ID (EPI_ISL_) | Amino acid change in S protein |
|---|---|---|---|
| WK-521 | JP/TY-WK-521/2020 | 408667 | - |
| SUMS2 | Japan/Shiga/SUMS2/2020 | 10434280 | D614G, Q675H |
| QHN-001 | Japan/QHN001/2020 | 804007 | deletion 69-70, deletion 144, N501Y, A570D, D614G, P681H, T716I, S982A, D1118H |
| TY7-501 | Japan/QHN001/2020 | 833366 | L18F, T20N, P26S, D138Y, G181V, R190S, K417T, E484K, N501Y, D614G, H655Y, T1027I, V1176F |
| TY8-612 | Japan/TY8-612/2021 | 1123289 | D80A, D215G, deletion, 242-244, K417N, E484K, N501Y, D614G, A701Y |
| TY11-927 | Japan/TY11-927-P1/2021 | 2158617 | T19R, G142D, E156G, deletion 157-158, L452R, T478K, D614G, P681R, D950N, |
| TY38-873 | Japan/TY38-873/2021 | 7418017 | A67V, deletion 69-70, T95I, G142D, deletion 143-145, deletion 211, L212I, insertion 214 EPE, G339D, S371L, S373P, S375F, K417N, N440K, G446S, S477N, T478K, E484A, Q493R, G496S, Q498R, N501Y, Y505H, T547K, D614G, H655Y, N679K, P681H, N764K, D796Y, N856K, Q954H, N969K, L981F, |

Strain names, abbreviations, GISAID IDs, and amino acid changes in the S protein of the variants are listed.

Supplemental Fig. 1

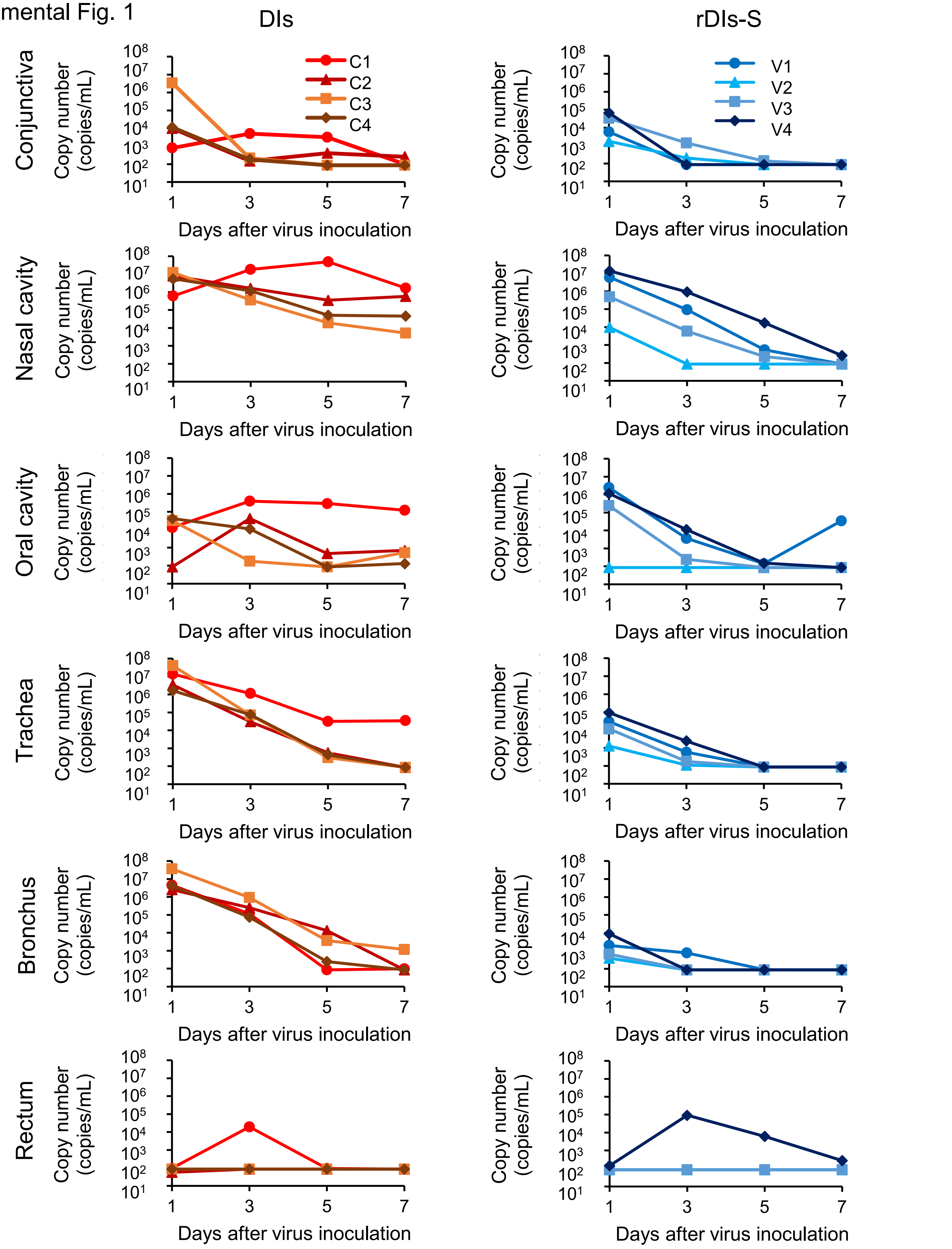


**Supplemental Figure 1. Viral RNA in swab samples of macaques vaccinated with rDIs-S.** Cynomolgus macaques were immunized intradermally with DIs (C1 – C4) or rDIs-S (V1 – V4). One week after the 2nd vaccination, WK-521 was inoculated into the conjunctiva, nostril, oral cavity, and trachea of each macaque on Day 0. Conjunctival, nasal, oral, tracheal, bronchial, and rectal swab samples then were collected on the indicated days. Viral RNA was quantified by qRT-PCR.

Supplemental Fig. 2

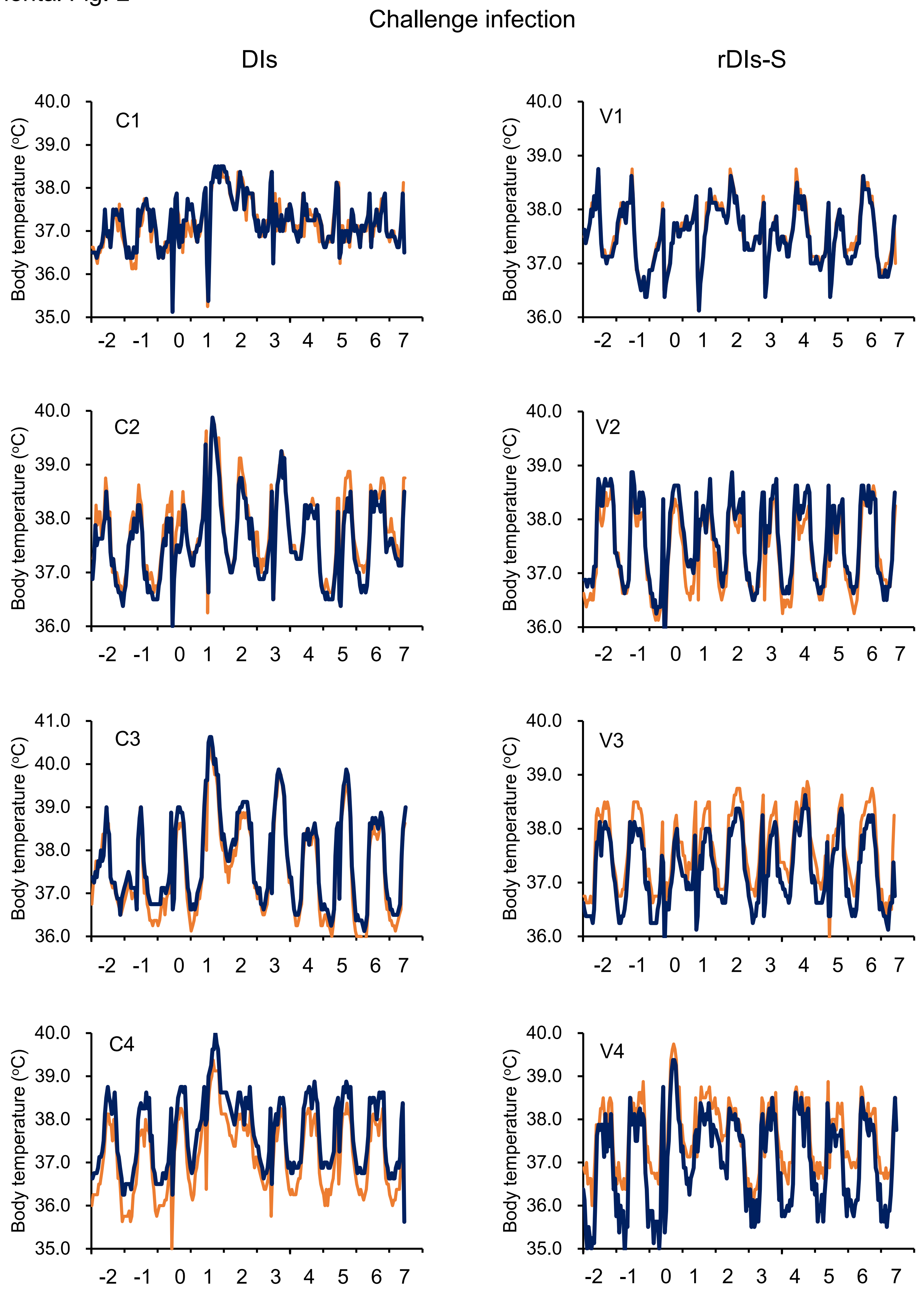


**Supplemental Figure 2. Body temperature of macaques after challenge infection with SARS-CoV-2.** The body temperatures of macaques immunized with DIs (C1 – C4) or rDIs-S (V1- V4) were recorded by two data loggers in each macaque (blue and orange lines). Animals were inoculated with SARS-CoV-2 WK-521 on Day 0.

Supplemental Fig. 3

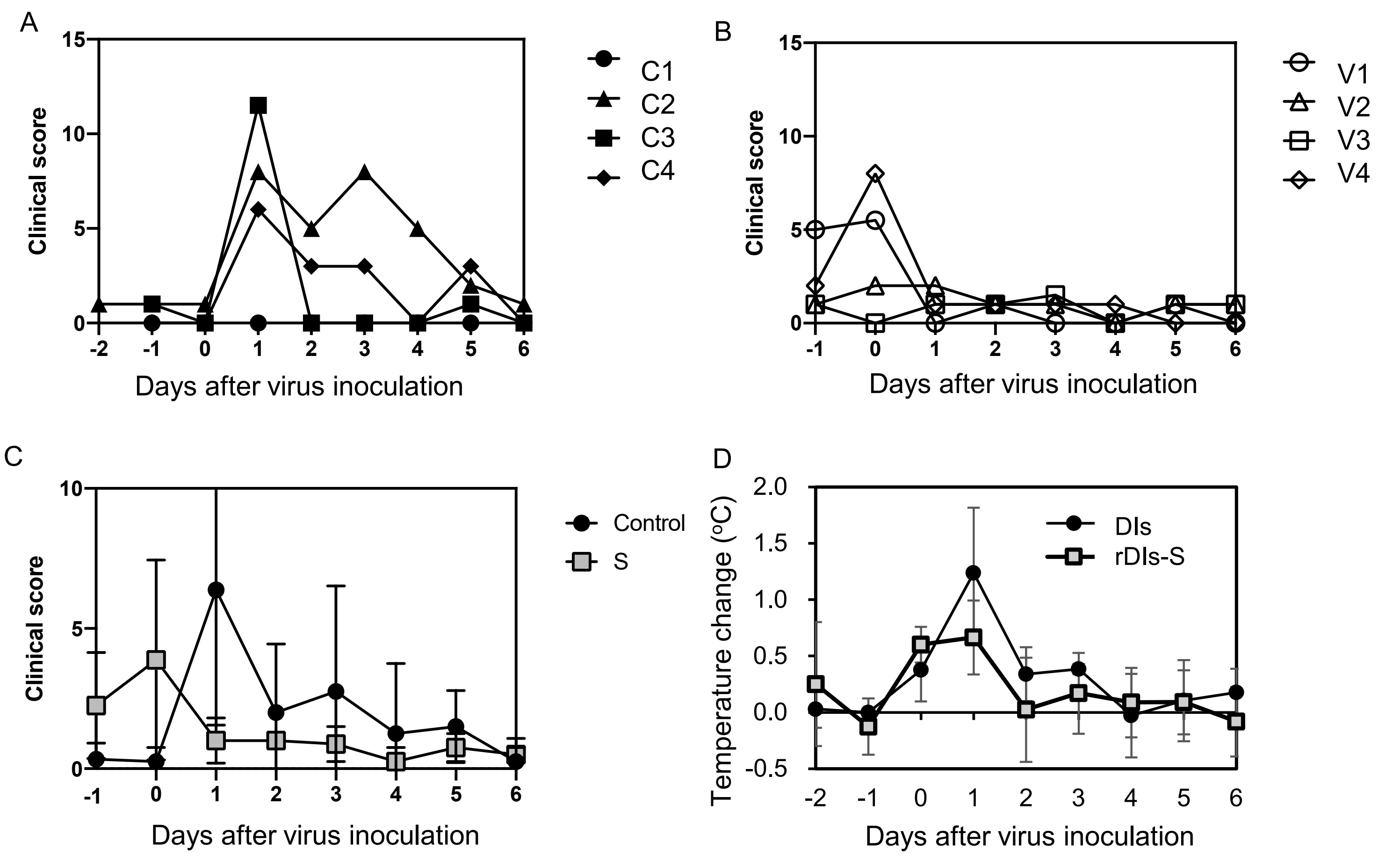


**Supplemental Figure 3. Clinical scores and mean body temperature change for macaques after challenge infection.**

(A, B) Macaques were observed every day and scored according to Supplemental Table 3. A: Macaques immunized with DIs. B: Macaques immunized with rDIs-S. (C) Mean and standard deviation of the clinical scores. (D) Mean body temperature changes. Means of the body temperatures of individual macaques from 8 p.m. to 8 a.m. were calculated each day. Body temperatures of the macaques were recorded using data loggers as indicated in legend for Supplemental Fig. 2. Mean body temperatures from 8 p.m. to 8 a.m. of the following day were calculated using the data in individual macaques, given that temperatures during the daytime were affected by anesthesia. For example, “the temperatures on Day 0” means the mean temperatures between 8 p.m. on Day 0 and 8 a.m. on Day 1 after virus inoculation. The mean body temperatures on each day were compared to those on Day -1 (from 8 p.m. on Day -1 to 8 a.m. on Day 0, before virus inoculation).

Supplemental Fig. 4

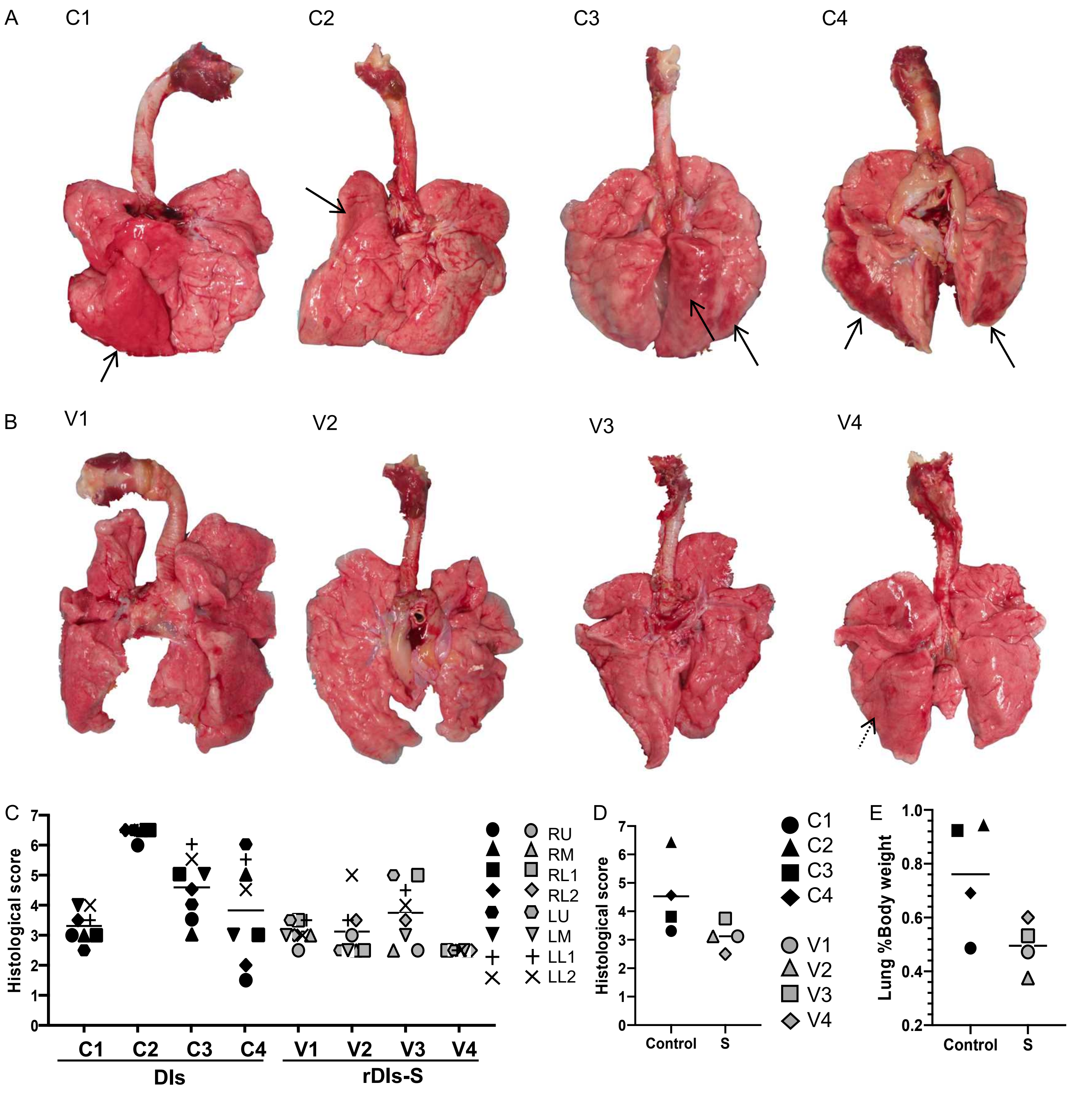


**Supplemental Figure 4. Gross appearance of lungs and histological scores of pneumonia.** (A, B) Gross appearance of lungs 7 days after virus inoculation. (A) Macaques immunized with DIs. (B) Macaques immunized with rDIs-S. Arrows: reddish lesions indicating pneumonia and congestion. (C) Histological scores of individual lung lobes of macaques 7 days after virus inoculation. Histological scores were determined independently by two pathologists (blinded to sample identity) based on the criteria described in the Methods. (D) Mean lung inflammation scores were calculated based on the results provided in Supplemental Fig. 4C. Bars indicate the mean histological scores. (E) Relative (body weight-normalized) lung weight is presented as percentages (= 100 × lung weight / body weight). Bars indicate the mean lung weight percentages.

Supplemental Fig. 5

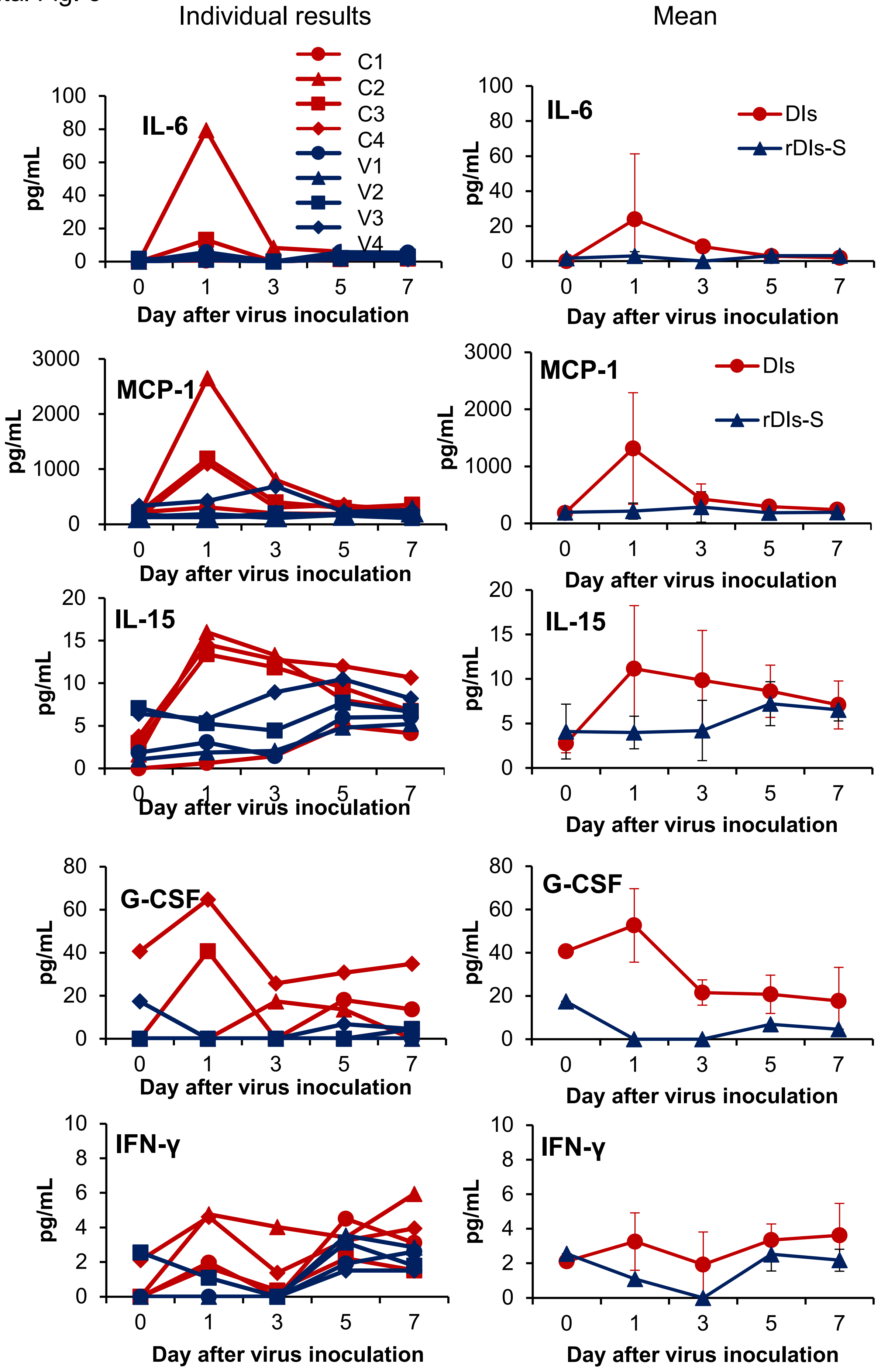


**Supplemental Figure 5. Plasma cytokine responses of macaques after challenge infection with SARS-CoV-2.** Plasma was collected on the indicated days after the challenge infection. The cytokine levels of individual macaques (left), and the means and standard deviations of four macaques (right) are shown. IL-6: interleukin-6; MCP-1: Monocyte chemoattractant protein-1; IL-15: interleukin-15; G-CSF: granulocyte colony-stimulating factor; IFN-γ: interferon-gamma.

Supplemental Fig. 6

A

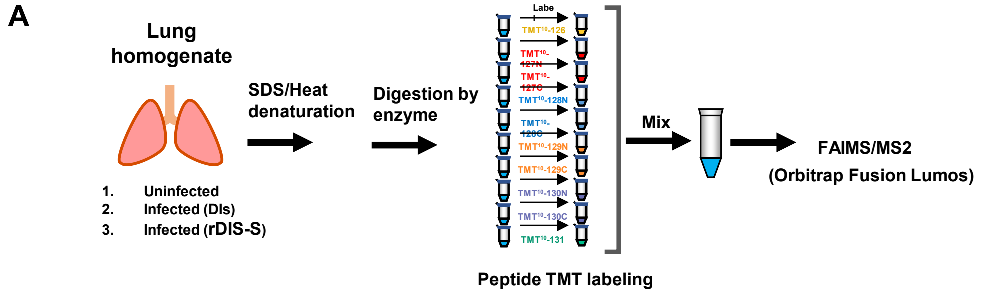


B

| Gene | Log2 FC | Log2 FC |
|---|---|---|
| Aldob | 2.56 | -0.32 |
| Arg1 | 1.87 | -0.30 |
| Bhmt | 3.50 | -0.28 |
| Camp | 2.80 | 0.32 |
| Chi3l3 | 2.02 | -0.26 |
| Elane | 2.15 | 0.15 |
| Fbp1 | 2.51 | 0.14 |
| Fga | 1.35 | -0.06 |
| Fgb | 1.20 | -0.02 |
| Fgg | 1.29 | 0.07 |
| Gnmt | 1.85 | 0.51 |
| Gp1ba | 1.19 | -0.20 |
| Hpd | 2.38 | -0.26 |
| Il1r2 | 3.35 | 0.14 |
| Itgam | 1.60 | 0.24 |
| Itgb2 | 1.26 | 0.03 |
| Itgb2l | 2.80 | 0.38 |
| Itih3 | 2.58 | 0.84 |
| Lpl | 2.23 | 0.04 |
| Ltf | 2.88 | 0.21 |
| Mmp8 | 2.85 | 0.40 |
| Mmp9 | 2.63 | 0.47 |
| Mpo | 2.25 | -0.13 |
| Mt2 | 1.53 | -0.44 |
| Ngp | 3.13 | 0.18 |
| Pck1 | 1.64 | -0.05 |
| Pglyrp1 | 3.15 | 0.22 |
| Plek | 1.57 | 0.23 |
| Prtn3 | 2.61 | -0.09 |
| S100a8 | 2.99 | 0.43 |
| S100a9 | 2.89 | 0.35 |
| Serinc3 | 1.70 | 0.34 |
| Serpina3c | 2.35 | 0.71 |
| Serpina3n | 1.27 | -0.21 |
| Stfa1 | 3.32 | 0.71 |
| Stfa2 | 2.97 | 0.89 |
| Tbxas1 | 1.74 | 0.18 |
| Thbs1 | 1.47 | -0.34 |
| Timp3 | 1.21 | -0.16 |

Log2 FC: 4, 2, 0, -2, -4

C

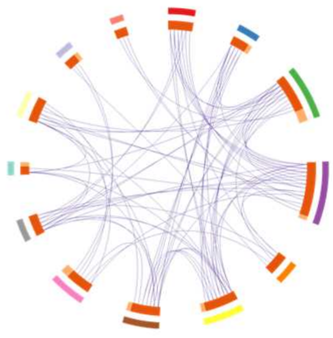

| GO enrichment terms | Number of proteins |
|---|---|
| leukocyte migration involved in inflammatory response | 6 |
| fibrinolysis | 5 |
| regulation of peptidase activity | 12 |
| inflammatory response | 14 |
| defense response to fungus | 5 |
| regulation of apoptotic signaling pathway | 9 |
| response to metal ion | 8 |
| small molecule biosynthetic process | 8 |
| positive regulation of angiogenesis | 5 |
| collagen catabolic process | 3 |
| response to peptide | 6 |
| alpha-amino acid metabolic process | 4 |
| aminoglycan metabolic process | 3 |

D

**Leukocyte migration involved in inflammatory response**

| Gene | Log2 (DIs/Ctrl.) | Log2 (rDIs-S/Ctrl.) |
|---|---|---|
| Itgam | 1.60 | 0.24 |
| Itgb2 | 1.26 | 0.03 |
| Itgb2l | 2.80 | 0.38 |
| S100a8 | 2.99 | 0.43 |
| S100a9 | 2.89 | 0.35 |
| Elane | 2.15 | 0.15 |

**Regulation of peptidase activity**

| Gene | Log2 (DIs/Ctrl.) | Log2 (rDIs-S/Ctrl.) |
|---|---|---|
| Itih3 | 2.58 | 0.84 |
| Serpina3c | 2.35 | 0.71 |
| Ltf | 2.88 | 0.21 |
| Mmp9 | 2.63 | 0.47 |
| Ngp | 3.13 | 0.18 |
| S100a8 | 2.99 | 0.43 |
| S100a9 | 2.89 | 0.35 |
| Serpina3n | 1.27 | -0.21 |
| Stfa1 | 3.32 | 0.71 |
| Stfa2 | 2.97 | 0.89 |
| Thbs1 | 1.47 | -0.34 |
| Timp3 | 1.21 | -0.16 |

**Defense response to fungus**

| Gene | Log2 (DIs/Ctrl.) | Log2 (rDIs-S/Ctrl.) |
|---|---|---|
| Arg1 | 1.87 | -0.30 |
| Camp | 2.80 | 0.32 |
| Ltf | 2.88 | 0.21 |
| Mpo | 2.25 | -0.13 |
| Elane | 2.15 | 0.15 |

**Regulation of apoptotic signaling pathway**

| Gene | Log2 (DIs/Ctrl.) | Log2 (rDIs-S/Ctrl.) |
|---|---|---|
| Fga | 1.35 | -0.06 |
| Mmp9 | 2.63 | 0.47 |
| S100a8 | 2.99 | 0.43 |
| S100a9 | 2.89 | 0.35 |
| Thbs1 | 1.47 | -0.34 |
| Timp3 | 1.21 | -0.16 |
| Serinc3 | 1.70 | 0.34 |
| Fgg | 1.29 | 0.07 |
| Fgb | 1.20 | -0.02 |

**Response to metal ion**

| Gene | Log2 (DIs/Ctrl.) | Log2 (rDIs-S/Ctrl.) |
|---|---|---|
| Fbp1 | 2.51 | 0.14 |
| Fga | 1.35 | -0.06 |
| Mmp9 | 2.63 | 0.47 |
| Mt2 | 1.53 | -0.44 |
| Thbs1 | 1.47 | -0.34 |
| Fgg | 1.29 | 0.07 |
| Fgb | 1.20 | -0.02 |
| Aldob | 2.56 | -0.32 |

**Small molecule biosynthetic process**

| Gene | Log2 (DIs/Ctrl.) | Log2 (rDIs-S/Ctrl.) |
|---|---|---|
| Bhmt | 3.50 | -0.28 |
| Fbp1 | 2.51 | 0.14 |
| Gnmt | 1.85 | 0.51 |
| Lpl | 2.23 | 0.04 |
| Mpo | 2.25 | -0.13 |
| Pck1 | 1.64 | -0.05 |
| Tbxas1 | 1.74 | 0.18 |
| Plek | 1.57 | 0.23 |

**Positive regulation of angiogenesis**

| Gene | Log2 (DIs/Ctrl.) | Log2 (rDIs-S/Ctrl.) |
|---|---|---|
| Camp | 2.80 | 0.32 |
| Itgb2 | 1.26 | 0.03 |
| Itgb2l | 2.80 | 0.38 |
| Mmp9 | 2.63 | 0.47 |
| Thbs1 | 1.47 | -0.34 |

**Response to peptide**

| Gene | Log2 (DIs/Ctrl.) | Log2 (rDIs-S/Ctrl.) |
|---|---|---|
| Serpina3c | 2.35 | 0.71 |
| Lpl | 2.23 | 0.04 |
| Mmp9 | 2.63 | 0.47 |
| Pck1 | 1.64 | -0.05 |
| Serpina3n | 1.27 | -0.21 |
| Aldob | 2.56 | -0.32 |

**Alpha-amino acid metabolic process**

| Gene | Log2 (DIs/Ctrl.) | Log2 (rDIs-S/Ctrl.) |
|---|---|---|
| Arg1 | 1.87 | -0.30 |
| Bhmt | 3.50 | -0.28 |
| Gnmt | 1.85 | 0.51 |
| Hpd | 2.38 | -0.26 |

**Aminoglycan metabolic process**

| Gene | Log2 (DIs/Ctrl.) | Log2 (rDIs-S/Ctrl.) |
|---|---|---|
| Chil3 | 2.02 | -0.26 |
| Itih3 | 2.58 | 0.84 |
| Pglyrp1 | 3.15 | 0.22 |

**Supplemental Figure 6. Relative quantitative analysis of proteins in lung tissues of hACE2 transgenic mice and GO term enrichment of terms related to BP of the upregulated proteins in the DIs-immunized mice.** (A) Summary of the procedures for the MS experiments with TMT Isobaric Mass Tagging Reagents. The protein extracts isolated from lung homogenates were reduced, alkylated, and digested. Samples were labeled with the TMT reagents, then pooled before sample fractionation and clean-up. Labeled samples were analyzed using a high-resolution Orbitrap Fusion Lumos (liquid chromatography-MS/MS). (B) List of genes encoding the proteins involved in the top-13 gene ontology (GO) enrichment terms related to biological processes (BP) among the upregulated proteins in the infected mice without vaccination when compared to the uninfected mice. (C) Overlap analysis of all proteins listed in Supplemental Fig. 6B. Outer arcs represent the GO enrichment terms indicated in the table on the right; the different groups are indicated by the color codes in the table. Inner arcs represent proteins in the protein lists (D), where each protein is indicated as a spot on the arc. Dark orange represents the proteins that appear in multiple lists, while light orange represents the proteins that are unique to that protein list. Purple lines link the same proteins that are shared between multiple protein lists. (D) Protein lists in residual clusters of the top-13 GO enrichment terms related to BP, except for the three clusters listed in Fig. 5E.

Supplemental Fig. 7

**A**

| | Infected (DIs)/Ctrl. | Infected (rDIs-S)/Ctrl. |
|---|---|---|
| Ahr | -1.25 | -0.12 |
| Cd74 | -1.19 | 0.16 |
| Cuedc2 | -1.76 | -0.26 |
| Cxcl15 | -2.74 | -0.27 |
| Ddr2 | -1.31 | -0.13 |
| Ear1 | -2.71 | -0.11 |
| Ear2 | -1.84 | -0.23 |
| Epx | -1.78 | 0.44 |
| H2-Aa | -1.57 | 0.06 |
| H2-Ab1 | -1.84 | -0.05 |
| Nrcam | -1.79 | -0.72 |
| Pcna | -1.48 | -0.45 |
| Pja2 | -2.04 | -0.83 |
| Prg2 | -2.47 | 0.96 |
| Twsg1 | -2.19 | -0.79 |
| Ybx1 | -1.85 | -0.82 |

Log2 FC: 4, 2, 0, -2, -4

**B**

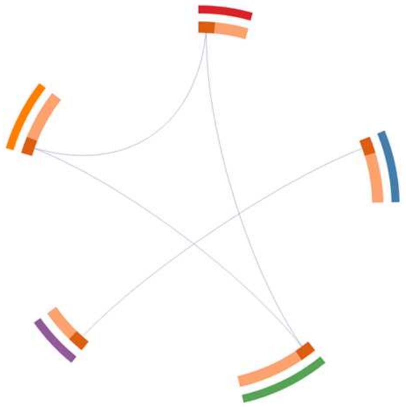

| GO enrichment terms | Number of proteins |
|---|---|
| antigen processing and presentation of exogenous peptide antigen via MHC class II | 3 |
| negative regulation of cytokine production | 4 |
| chemotaxis | 5 |
| regulation of osteoblast differentiation | 3 |
| regulation of cellular response to stress | 4 |

**C**

**Negative regulation of cytokine production**

| Protein | Log2 (DIs/Ctrl.) | Log2 (rDIs-S/Ctrl.) |
|---|---|---|
| Epx | -1.78 | 0.44 |
| Prg2 | -2.47 | 0.96 |
| Twsg1 | -2.19 | -0.79 |
| Cuedc2 | -1.76 | -0.26 |

**Chemotaxis**

| Protein | Log2 (DIs/Ctrl.) | Log2 (rDIs-S/Ctrl.) |
|---|---|---|
| Ear1 | -2.71 | -0.11 |
| Ear2 | -1.84 | -0.23 |
| Cd74 | -1.19 | 0.16 |
| Cxcl15 | -2.74 | -0.27 |
| Nrcam | -1.79 | -0.72 |

**Regulation of osteoblast differentiation**

| Protein | Log2 (DIs/Ctrl.) | Log2 (rDIs-S/Ctrl.) |
|---|---|---|
| Ahr | -1.25 | -0.12 |
| Ddr2 | -1.31 | -0.13 |
| Twsg1 | -2.19 | -0.79 |

**Supplemental Figure 7. GO enrichment terms related to BP of the downregulated proteins in the DIs-immunized mice.**

(A) The top-five gene ontology (GO) enrichment terms related to biological processes (BP) of proteins that were downregulated in the infected mice without vaccination when compared to the uninfected mice. (B) Overlap analysis of all proteins listed in Supplemental Fig. 7A. GO enrichment terms of proteins are indicated in the table on the right; the different groups are indicated by the color codes in the table. Inner arcs represent proteins in the protein lists (C), where each protein is indicated as a spot on the arc. Dark orange represents the proteins that appear in multiple lists, while light orange represents the proteins that are unique to that protein list. Purple lines link the same proteins that are shared by multiple protein lists. (C) Protein lists in residual clusters of the top-5 GO enrichment terms related to BP, except for the two clusters listed in Fig. 5F.

### Supplemental Fig. 8

**A** **Up-regulated proteins in both groups**

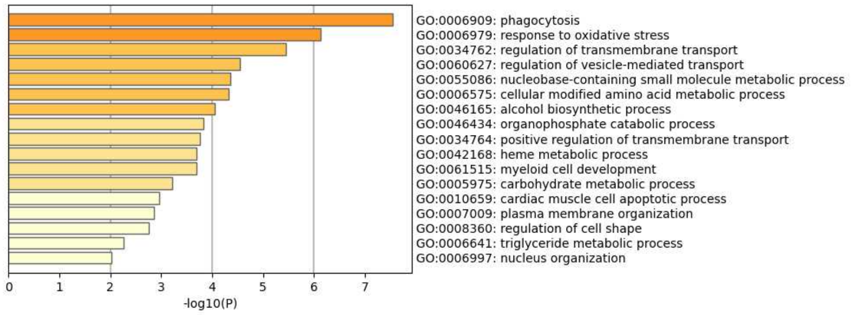


**B**

| | Infected (DIs)/Ctrl. | Infected (rDIs-S)/Ctrl. |
|---|---|---|
| Acsl1 | 1.11 | 1.23 |
| Apaf1 | 1.28 | 1.04 |
| Blvrb | 1.80 | 1.34 |
| Bpifa1 | 1.89 | 1.59 |
| Cdk5 | 1.02 | 1.10 |
| Cmas | 1.78 | 2.00 |
| Coq7 | 1.30 | 1.19 |
| Coro1a | 1.87 | 1.69 |
| Cpt1b | 1.22 | 1.18 |
| Dhfr | 1.55 | 1.39 |
| Eef1g | 2.01 | 2.00 |
| Epb41l3 | 1.48 | 1.31 |
| Fes | 1.95 | 1.26 |
| G6pdx | 1.89 | 1.43 |
| Gclm | 1.37 | 1.51 |
| Gpcpd1 | 1.62 | 1.29 |
| Gpld1 | 1.47 | 1.10 |
| Hk3 | 1.90 | 1.17 |
| Hmbs | 1.46 | 1.72 |
| Hmgcs2 | 2.32 | 1.66 |
| Ighg1 | 3.40 | 1.77 |
| Ighg2a | 2.97 | 2.12 |
| Ighg2b | 2.47 | 1.55 |
| Ighv1-74 | 2.60 | 1.55 |
| Ighv3-6 | 2.71 | 1.81 |
| Ighv5-17 | 2.81 | 2.14 |
| Igkc | 2.49 | 1.48 |
| Mapk14 | 1.67 | 1.54 |
| Mlycd | 2.00 | 1.95 |
| Nt5c3 | 2.22 | 1.56 |
| Osbpl2 | 2.02 | 1.74 |
| Pcmt1 | 1.18 | 1.24 |
| Plcg2 | 1.80 | 1.18 |
| Pnkp | 1.70 | 1.40 |
| Psme4 | 1.25 | 1.17 |
| Pten | 1.10 | 1.53 |
| Ptpn11 | 1.04 | 1.13 |
| Rhd | 1.41 | 1.02 |
| Slc43a1 | 1.38 | 1.02 |
| Slc4a1 | 1.30 | 1.12 |
| Sphk1 | 2.06 | 1.38 |
| Thg1l | 1.10 | 1.14 |
| Tor1a | 1.22 | 1.16 |
| Urod | 1.19 | 1.03 |
| Zmpste24 | 1.25 | 1.16 |

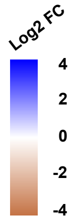


**C**

Phagocytosis

| Protein | Log2 (DIs/Ctrl.) | Log2 (rDIs-S/Ctrl.) |
|---|---|---|
| Coro1a | 1.87 | 1.69 |
| Ighg2b | 2.47 | 1.55 |
| Ighg1 | 3.40 | 1.77 |
| Igkc | 2.49 | 1.48 |
| Pten | 1.10 | 1.53 |
| Sphk1 | 2.06 | 1.38 |
| Plcg2 | 1.80 | 1.18 |
| Ighg2a | 2.97 | 2.12 |
| Ighv5-17 | 2.81 | 2.14 |
| Ighv3-6 | 2.71 | 1.81 |
| Ighv1-74 | 2.60 | 1.55 |

Response to oxidative stress

| Protein | Log2 (DIs/Ctrl.) | Log2 (rDIs-S/Ctrl.) |
|---|---|---|
| Coq7 | 1.30 | 1.19 |
| Dhfr | 1.55 | 1.39 |
| G6pdx | 1.89 | 1.43 |
| Gclm | 1.37 | 1.51 |
| Slc4a1 | 1.30 | 1.12 |
| Sphk1 | 2.06 | 1.38 |
| Tor1a | 1.22 | 1.16 |
| Pnkp | 1.70 | 1.40 |
| Thg1l | 1.10 | 1.14 |
| Hk3 | 1.90 | 1.17 |

Regulation of transmembrane transport

| Protein | Log2 (DIs/Ctrl.) | Log2 (rDIs-S/Ctrl.) |
|---|---|---|
| Cdk5 | 1.02 | 1.10 |
| Coro1a | 1.87 | 1.69 |
| Acsl1 | 1.11 | 1.23 |
| G6pdx | 1.89 | 1.43 |
| Bpifa1 | 1.89 | 1.59 |
| Pten | 1.10 | 1.53 |
| Ptpn11 | 1.04 | 1.13 |
| Mapk14 | 1.67 | 1.54 |
| Slc43a1 | 1.38 | 1.02 |
| Zmpste24 | 1.25 | 1.16 |
| Plcg2 | 1.80 | 1.18 |

Regulation of vesicle-mediated transport

| Protein | Log2 (DIs/Ctrl.) | Log2 (rDIs-S/Ctrl.) |
|---|---|---|
| Cdk5 | 1.02 | 1.10 |
| Coro1a | 1.87 | 1.69 |
| Fes | 1.95 | 1.26 |
| Ighg2b | 2.47 | 1.55 |
| Ighg1 | 3.40 | 1.77 |
| Pten | 1.10 | 1.53 |
| Sphk1 | 2.06 | 1.38 |
| Tor1a | 1.22 | 1.16 |
| Plcg2 | 1.80 | 1.18 |
| Ighg2a | 2.97 | 2.12 |

Nucleobase-containing small molecule metabolic process

| Protein | Log2 (DIs/Ctrl.) | Log2 (rDIs-S/Ctrl.) |
|---|---|---|
| Cmas | 1.78 | 2.00 |
| Acsl1 | 1.11 | 1.23 |
| G6pdx | 1.89 | 1.43 |
| Hmgcs2 | 2.32 | 1.66 |
| Pcmt1 | 1.18 | 1.24 |
| Slc4a1 | 1.30 | 1.12 |
| Mlycd | 2.00 | 1.95 |
| Nt5c3 | 2.22 | 1.56 |
| Hk3 | 1.90 | 1.17 |

Cellular modified amino acid metabolic process

| Protein | Log2 (DIs/Ctrl.) | Log2 (rDIs-S/Ctrl.) |
|---|---|---|
| Cpt1b | 1.22 | 1.18 |
| Dhfr | 1.55 | 1.39 |
| G6pdx | 1.89 | 1.43 |
| Gclm | 1.37 | 1.51 |
| Pcmt1 | 1.18 | 1.24 |
| Eef1g | 2.01 | 2.00 |

Alcohol biosynthetic process

| Protein | Log2 (DIs/Ctrl.) | Log2 (rDIs-S/Ctrl.) |
|---|---|---|
| Dhfr | 1.55 | 1.39 |
| G6pdx | 1.89 | 1.43 |
| Hmgcs2 | 2.32 | 1.66 |
| Sphk1 | 2.06 | 1.38 |
| Plcg2 | 1.80 | 1.18 |

Organophosphate catabolic process

| Protein | Log2 (DIs/Ctrl.) | Log2 (rDIs-S/Ctrl.) |
|---|---|---|
| Pten | 1.10 | 1.53 |
| Mlycd | 2.00 | 1.95 |
| Gpcpd1 | 1.62 | 1.29 |
| Nt5c3 | 2.22 | 1.56 |
| Plcg2 | 1.80 | 1.18 |

Positive regulation of transmembrane transport

| Protein | Log2 (DIs/Ctrl.) | Log2 (rDIs-S/Ctrl.) |
|---|---|---|
| Cdk5 | 1.02 | 1.10 |
| Acsl1 | 1.11 | 1.23 |
| G6pdx | 1.89 | 1.43 |
| Ptpn11 | 1.04 | 1.13 |
| Mapk14 | 1.67 | 1.54 |
| Plcg2 | 1.80 | 1.18 |

Heme metabolic process

| Protein | Log2 (DIs/Ctrl.) | Log2 (rDIs-S/Ctrl.) |
|---|---|---|
| Hmbs | 1.46 | 1.72 |
| Urod | 1.19 | 1.03 |
| Blvrb | 1.80 | 1.34 |

**Supplemental Figure 8. GO BP enrichment terms of the upregulated proteins in both infected groups.** (A) The top-17 gene ontology (GO) enrichment terms related to biological processes (BP) of the upregulated proteins in the infected groups (both the control (DIs) and experimental (rDIs-S) vaccinated groups). (B) List of proteins involved in the top-10 GO enrichment terms related to BP of the upregulated proteins in both infected groups compared to the uninfected group. (C) Protein lists of the clusters of the top-10 GO enrichment terms related to BP.

Supplemental Fig. 9

**A Down-regulated proteins in both groups**

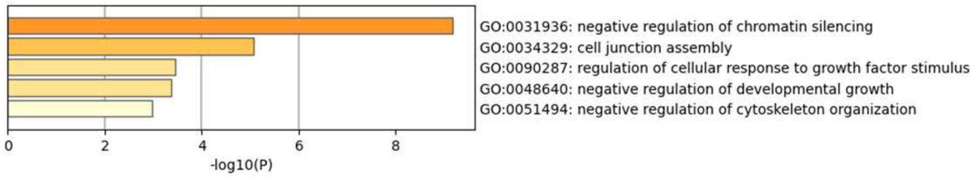


**B**

| | Infected (DIs)/Ctrl. | Infected (rDIs-S)/Ctrl. |
|---|---|---|
| Apc | -1.20 | -1.06 |
| Cgnl1 | -1.47 | -1.09 |
| Epn2 | -1.31 | -1.19 |
| Fstl1 | -2.37 | -1.02 |
| Gap43 | -1.26 | -1.13 |
| H1f1 | -1.38 | -1.54 |
| H1f3 | -1.61 | -1.75 |
| H1f4 | -1.57 | -1.57 |
| H1f5 | -2.94 | -3.12 |
| Jcad | -1.10 | -1.12 |
| Kdr | -1.54 | -1.07 |
| Ntn1 | -1.13 | -1.06 |
| Peak1 | -1.30 | -1.01 |
| Phldb2 | -1.25 | -1.12 |

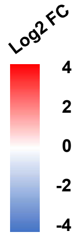


**C**

**Negative regulation of chromatin silencing**

| Protein | Log2 (DIs/Ctrl.) | Log2 (rDIs-S/Ctrl.) |
|---|---|---|
| H1f3 | -1.61 | -1.75 |
| H1f4 | -1.57 | -1.57 |
| H1f5 | -2.94 | -3.12 |
| H1f1 | -1.38 | -1.54 |

**Regulation of cell-matrix adhesion**

| Protein | Log2 (DIs/Ctrl.) | Log2 (rDIs-S/Ctrl.) |
|---|---|---|
| Kdr | -1.54 | -1.07 |
| Phldb2 | -1.25 | -1.12 |
| Peak1 | -1.30 | -1.01 |

**Response to growth factor**

| Protein | Log2 (DIs/Ctrl.) | Log2 (rDIs-S/Ctrl.) |
|---|---|---|
| Epn2 | -1.31 | -1.19 |
| Fstl1 | -2.37 | -1.02 |
| Kdr | -1.54 | -1.07 |
| Jcad | -1.10 | -1.12 |

**Negative regulation of organelle organization**

| Protein | Log2 (DIs/Ctrl.) | Log2 (rDIs-S/Ctrl.) |
|---|---|---|
| Apc | -1.20 | -1.06 |
| Cgnl1 | -1.47 | -1.09 |
| Phldb2 | -1.25 | -1.12 |

**Cell part morphogenesis**

| Protein | Log2 (DIs/Ctrl.) | Log2 (rDIs-S/Ctrl.) |
|---|---|---|
| Apc | -1.20 | -1.06 |
| Gap43 | -1.26 | -1.13 |
| Kdr | -1.54 | -1.07 |
| Ntn1 | -1.13 | -1.06 |

**Supplemental Figure 9. GO BP enrichment terms of the downregulated proteins in both infected groups.**

Top-five gene ontology (GO) enrichment terms related to biological processes (BP) of the downregulated proteins in the infected groups (both the control (DIs) and experimental (rDIs-S) vaccinated groups). (B) List of proteins involved in the top-five GO enrichment terms related to BP of the downregulated proteins in both infected groups compared to the uninfected group. (C) Protein lists of the clusters of the top-five GO enrichment terms related to BP.
